# Codon-dependent noise dictates cell-to-cell variability in nutrient poor environments

**DOI:** 10.1101/492207

**Authors:** Enrique Balleza, Lisa F. Marshall, J. Mark Kim, Philippe Cluzel

## Abstract

Under nutrient-rich conditions, stochasticity of transcription drives protein expression noise. However, by shifting the environment to amino acid-limited conditions, we identified in E. coli a source of noise whose strength is dictated by translational processes. Specifically, we discovered that cell-to-cell variations in fluorescent protein expression depend on codon choice, with codons yielding lower mean expression after amino acid downshift also resulting in greater noise. We propose that ultra-sensitivity in the tRNA charging/discharging cycle shapes the strength of the observed noise by amplifying fluctuations in global intracellular parameters, such as the concentrations of amino acid, synthetase, and tRNA. We hypothesize that this codon-dependent noise may allow bacteria to selectively optimize cell-to-cell variability in poor environments without relying on low molecular numbers.

---

Genetically identical cells exposed to homogeneous environments can nonetheless exhibit dramatic variations in gene expression, commonly termed “noise” (*1, 2*). High quality single cell measurements combined with mathematical modeling has led to a quantitative understanding of the main sources of noise in gene expression (*3, 4*). In bacteria, these studies have established that that noise primarily stems from transcription (*3*–*5*) not translation. However, most of these studies were performed in nutrient rich environments where translation is not the limiting process in protein synthesis—by contrast, very little is known about noise when bacteria grow in nutrient poor environments. To tackle this problem in *E. coli*, we built on the approach developed in (*6*), which demonstrated that the choice of certain synonymous codons could govern protein levels when bacteria starve for the cognate amino acid. For example, prompted by computational predictions (*7*), we found that codons encoding leucine can be split into two categories: those that are more robust and those that are more sensitive to leucine starvation. Under leucine-limited conditions, we showed that levels of the yellow fluorescent protein (YFP) expression are much greater when expressed from gene sequences using the robust leucine codon CGT in place of the sensitive codon CTA. In the light of this earlier work, we decided to characterize how synonymous codon choice might affect not only the protein levels but also the associated noise when the growth environment is downshifted from amino acid-rich media (“rich” condition) to media limited for the cognate amino acid (“starvation” condition).

To quantify gene expression and noise levels, we engineered a strain with a chromosomal copy of *yfp* stably expressed from a Tet-inducible promoter (P_LTetO-1_). Expression of YFP was only induced at the time of nutrient downshift to avoid unwanted fluorescence from any YFP produced before starvation. We subsequently harvested the cells and estimated YFP expression levels and the associated noise using flow cytometry (**Supplementary Fig. 3**). Operationally, we used leucine codons as a testbed for characterizing noise during starvation, because the response to leucine starvation is faster and greater, which makes it more amenable to precise measurements. However, we extended this approach to other amino acids (**Supplementary Note**). In order to decouple the noise due to translation from that due to transcription after an amino acid downshift, we performed most of our experiments in the leucine auxotroph Δ*leuB*.

In rich media, the standard model for expression of unregulated genes assumes that translation does not significantly contribute to noise (Fig. 1a), because the protein number per transcript is large (*3*). Consequently in rich media, cell-to-cell variability in mRNA levels is tightly correlated with variations in the protein level. This assumption implies that the source of noise is purely transcriptional and that the coefficient of variation associated with protein levels would remain constant even if the synthesis rate of proteins changes. We checked this key hypothesis of the standard model by modifying in two distinct ways the translation rate, either by changing the initiation rate or by substituting synonymous codons.

**Figure 1.**
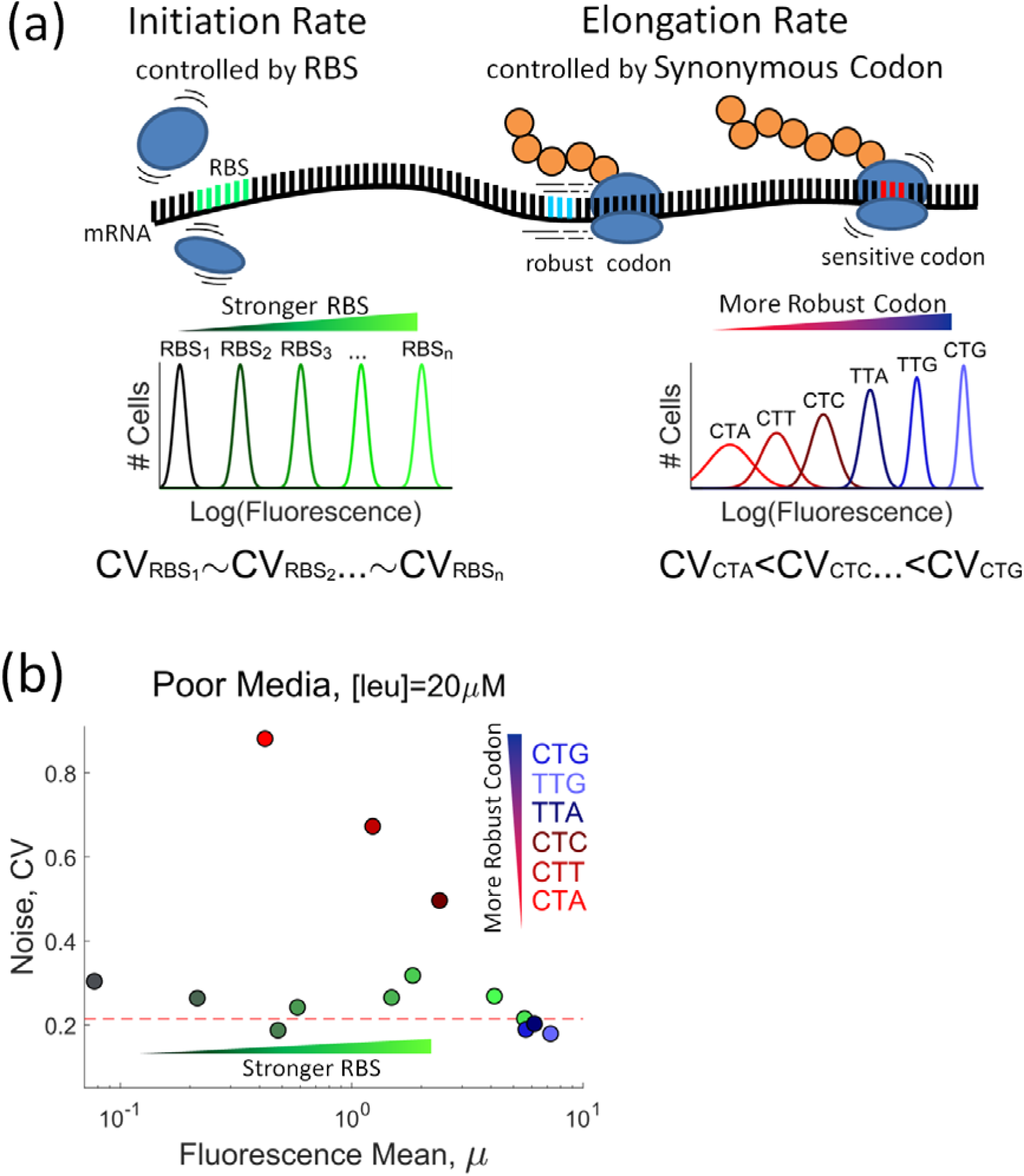
The choice of synonymous codons underlies noise in gene expression during leucine limitation. **(a)** Expectations on protein translation and associated noise during amino acid limitation when changing translation initiation rate via the ribosomal binding site (green), RBS, or when changing elongation rate via synonymous codons (sensitive in red, robust in blue). We represented, schematically, fluorescence distributions of cell populations expressing a YFP reporter. In the RBS case, distributions have the same width but change their mean according to the RBS strength, i.e. protein noise is roughly independent of initiation rate or, equivalently, noise comes mainly from transcription. For the synonymous codon case, the fluorescence distribution changes width and becomes tighter as more robust-to-starvation codons are used in the reporter sequence, i.e. depending on codon choice, the translation step can add a sizable amount of noise to protein expression. **(b)** Effect on noise of either different RBSs or different leucine synonymous codons in leucine-poor synthetic media, [leu]=20uM. Green circles: data from 8 different strains with RBSs of various strengths to modulate initiation rate. All RBSs drive the expression of a YFP that is coded with a robust-to-starvation codon, CTG. Red to blue circles: strains expressing YFP coded with one of the 6 possible synonymous leucine codons. Each strain uses an identical RBS to drive the expression of the synonymous variants. The dashed red line corresponds to the average noise level of the RBS library in leucine-rich synthetic media, [leu]=1mM. For each data point, we derived mean fluorescence values and CVs from at least 3×10^3^ cells. Error bars associated with the standard error of the mean are smaller than symbols sizes. We replicated the experiments in this figure using another set of independent RBS and leucine codon strains, see **Supplementary Fig. 5** and **7**.

First, we created a library of “robust” YFPs—i.e. YFPs coding sequences that use only robust codons as described in (*6*)—with ribosomal binding sites (RBS) of different strengths to modulate the translation initiation rate. In these constructs, a chromosomal copy of the *yfp* was stably expressed from an inducible Tet promoter and translation initiation rates were controlled by a series of ribosome binding sites of various strengths (**Supplementary Note**). Under rich growth conditions, and after full aTc induction, we found, as predicted by the standard model, that the noise (coefficient of variation) was unaffected by the strength of ribosomal binding sites (Fig. 1b). Next, we repeated the same experiment but transferred cells to a leucine-limited environment and found that the noise remained similar as in rich conditions, demonstrating that noise is independent from variations in translation initiation rates (*3, 4*) (**Supplementary Fig. 6-7**).

Second, we tested six different constructs of *yfp* re-coded with different synonymous leucine codons ranging from robust to sensitive. In these experiments, the initiation rate was kept constant by using only one specific ribosome binding site common to all constructs (**Supplementary Note**). Surprisingly, in leucine poor media (10uM range), the noise drastically deviated from the standard model: the construct using the most sensitive leucine codon, CTA, yielded more than a 4-fold noise increase than the construct with the robust codon, CTG (Fig. 1b). This novel source of noise, hereafter called ‘codon noise’, is not present in rich growth conditions where *yfp* expression noise associated with either robust and sensitive codons is not larger than the background transcription noise (Fig. 1b and **Supplementary Fig. 6**). Remarkably, however, in leucine poor media the amplitude of the codon noise follows the same hierarchy as hierarchy of mean expression levels found in (*6*): robust codons are quiet and sensitive codons are noisy under leucine starvation.

In order to understand the underlying mechanism that governs the magnitude of codon noise, we systematically measured how the expression levels (mean YFP fluorescence) and its associated noise (cell-to-cell variability) depends on leucine concentration. We measured for our six leucine constructs the mean fluorescence levels when cells where transferred from rich media to leucine poor medium ranging from 1mM down to 1uM. We found that the mean fluorescence decreased as a function of [leucine] in a sigmoidal fashion: constructs using the most sensitive codons (*i.e*. CTA, CTT, CTC) yielded the sharpest sigmoidal response as a function of leucine concentration (**Fig. 2 left**). By contrast, *yfp* genes using more robust codons (TTA, TTG, CTG) yielded shallower sigmoidal curves. Interestingly, the noise peaks in the region where the fluorescence curves exhibit a maximal *logarithmic gain* or *sensitivity amplification* (**Fig. 2 left, inset**) as previously predicted in (*7*), whereas robust codons yield only a modest noise increase, about a third of the noise observed in sensitive codons (**Fig. 2, right**).

**Figure 2.**
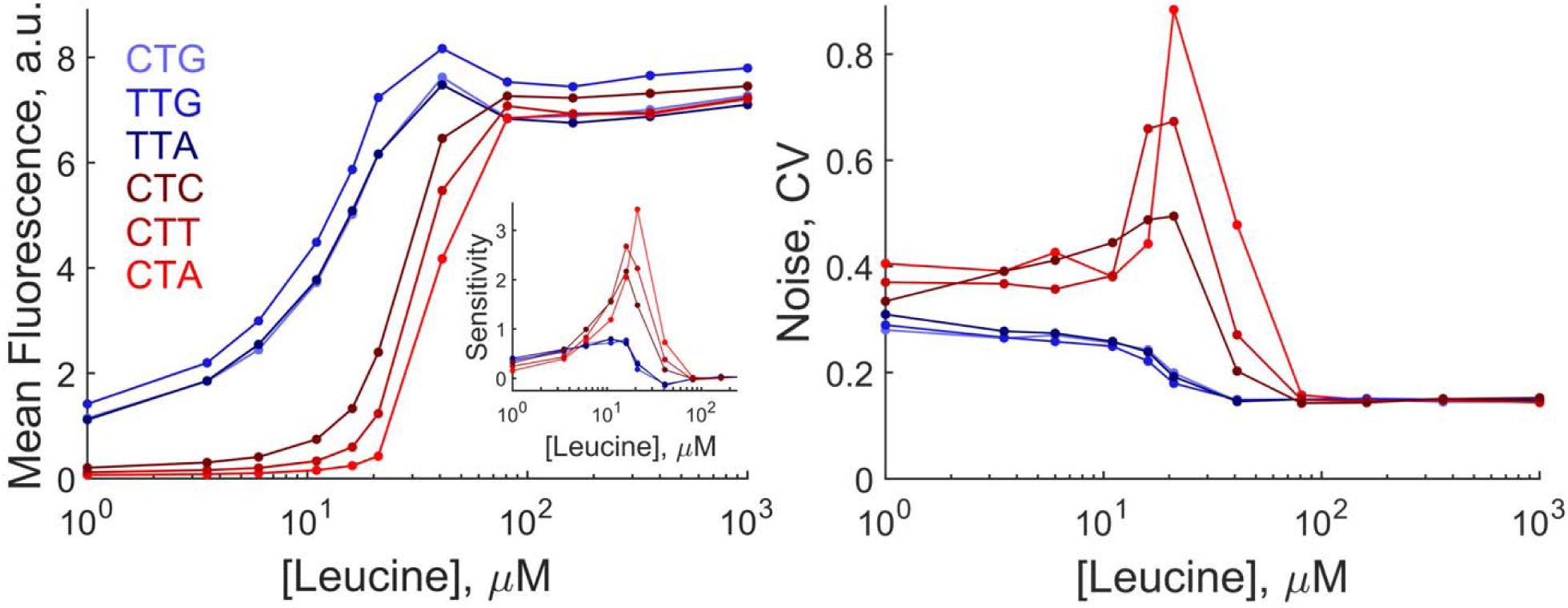
Robust and sensitive codons are quieter and noisier, respectively. Mean fluorescence (left) and associated noise (right) of a YFP reporter as a function of the concentration of leucine and the synonymous codon choice of the YFP reporter for the leucine codon family. We grew a Δ*leuB E. coli* strain (strict auxotroph in synthetic media without leucine) bearing a repressed YFP reporter in rich synthetic media until early exponential phase. Then, we transferred 1ul of culture into 1000ul of synthetic media with a predefined concentration of leucine and inducer (aTc) and incubated for 1hr. Finally, we transferred 240ul of culture into pre-chilled 96 microtiter wells and measured using a flow cytometer. The inset shows the sensitivity amplification or logarithmic gain of the mean fluorescence curves, (Δ*μ*/*μ*)/(Δ*Leu*/*Leu*). Note how high sensitivity amplification is associated to large random fluctuations in YFP synthesis (high CV). Mean and CV values were derived from a sample size of 3×10^3^ ± 200 cells. Error bars associated with the standard error of the mean are smaller than symbols sizes. We replicated the leucine concentration response and associated noise for another set of independent strains, see **Supplementary Fig. 5**.

While complex and intricate networks of interacting components are involved in protein synthesis, we nonetheless sought to discriminate intracellular parameters underlying the observed codon noise by adapting a well-known 2-color experiment developed by Elowitz *et al*. for analyzing transcription noise (*8*). We sought to identify whether codon noise is caused by processes that affect the translation machinery locally at the cognate codon in the transcript, or if codon noise is caused by cell-to-cell variability of more global intracellular parameters (e.g. synthetase or amino acid levels) (*3, 8*–*10*). To this end, we monitored the expression of a yellow and cyan fluorescent protein, *cfp and yfp*, controlled by two identical copies of the same ribosome binding sequence (**Supplementary Note**). We encoded the pair of the fluorescent proteins with sensitive leucine codons and simultaneously measured the fluorescence signal from CFP and YFP in individual cells via flow cytometry after a transfer from a rich to a leucine poor medium. A fluorescence scatterplot from flow cytometric measurements showed that CFP and YFP signals were strongly correlated along the diagonal and maximally spread over two orders of magnitude when the total codon noise peaked at [leucine]=20uM (Fig. 3a). The strong correlation along the diagonal demonstrates that codon noise is dominated by fluctuations of global intracellular parameters that affect both YFP and CFP transcripts: such global parameters include, but are not limited to, concentrations of the amino acid, synthetase or tRNA. In Figure 3b, we summarized for various leucine concentrations (1uM-1mM) the contribution of local and global noise to the total codon noise: as shown, global noise dominates codon noise throughout this leucine concentration range. That the **inset** of **Fig. 2 left** strongly correlates with the independent measurements of the coefficient of variation in **Fig. 2 right**, suggests that codon noise emerges from the sensitivity amplification of this system (*11*–*13*), which reflects the relative change of YFP synthesis rate normalized to the relative change in the amino acid supply. These latter results together with the fact that codon noise is dominated by fluctuations of global parameters led us to propose a minimal model.

**Figure 3.**
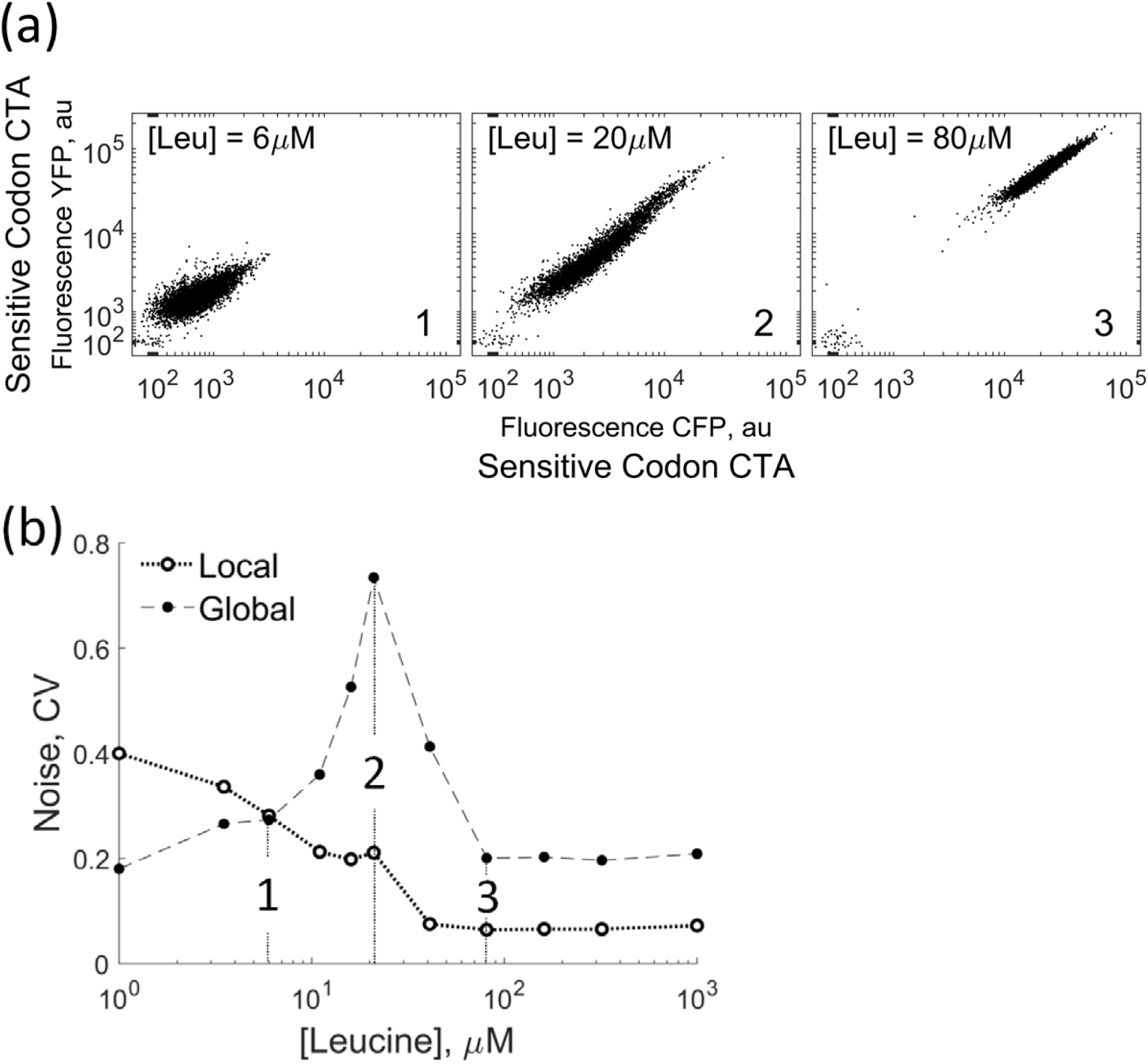
Fluctuations in global factors underlie translation noise. We constructed 2-color reporter strains to quantify the relative contribution of local and global factors to the observed translation noise at different amino acid concentrations. **(a)** Scattered plot from flow cytometry data at [leucine]=6uM, 20uM and 80uM of a 2-color reporter strain expressing yellow and cyan fluorescent proteins coded with the sensitive leucine codon CTA. **(b)** Noise decomposition into local and global components as a function of leucine concentration when both proteins are coded with CTA. Note that the scatter plots in **(a)** are labelled 1, 2, and 3 in **(b)**. The local and global CV values were derived from a sample size of 5×10^3^ ± 200 cells. Error bars associated with the standard error of the mean are smaller than symbols sizes. We replicated the experiments in this figure using another set of independent 2-color strains, see **Supplementary Fig. 9**.

First, in view of the complexity of translation processes in bacteria, we decided to simplify our theoretical model to isolate the essential controlling parameters of observed codon noise. We then considered a system where translation elongation rate is limited by a simple enzymatic cycle of charging-discharging of tRNA for one single cognate amino acid. In this cycle, the forward reaction corresponds to the tRNA amino-acylation reaction catalyzed by the cognate synthetase and the backward reaction is reduced to translation of a cognate codon. We modeled the forward and backward reactions with Michaelis-Menten kinetics (*14, 15*). In this framework, we used the currently available biochemical parameters to simulate the dependence of the concentration of charged tRNA (cognate to a sensitive codon) on the internal leucine concentration (**Supplementary Note**). Under this condition, we found that the synthetase is working at saturation, which yields the experimentally observed ultra-sensitive sigmoidal response of sensitive codons to [leucine] (Fig. 4b). Additionally, we found that one condition for which this minimal model would exhibit a peak within sharp sigmoidal transition was by assuming Poisson noise in the intracellular concentration of a global parameter such as amino acid concentration (Fig. 4a). But cell-to-cell variations in other global parameters could contribute to global noise as well. We interpret this peak as the results of amplifying property of the charging/discharging cycle of global noise. This computational result together with the observed peak of extrinsic noise in the 2-color experiment support the hypothesis that global parameters cell-to-cell variability dominate the observed codon noise (Fig. 2).

**Figure 4.**
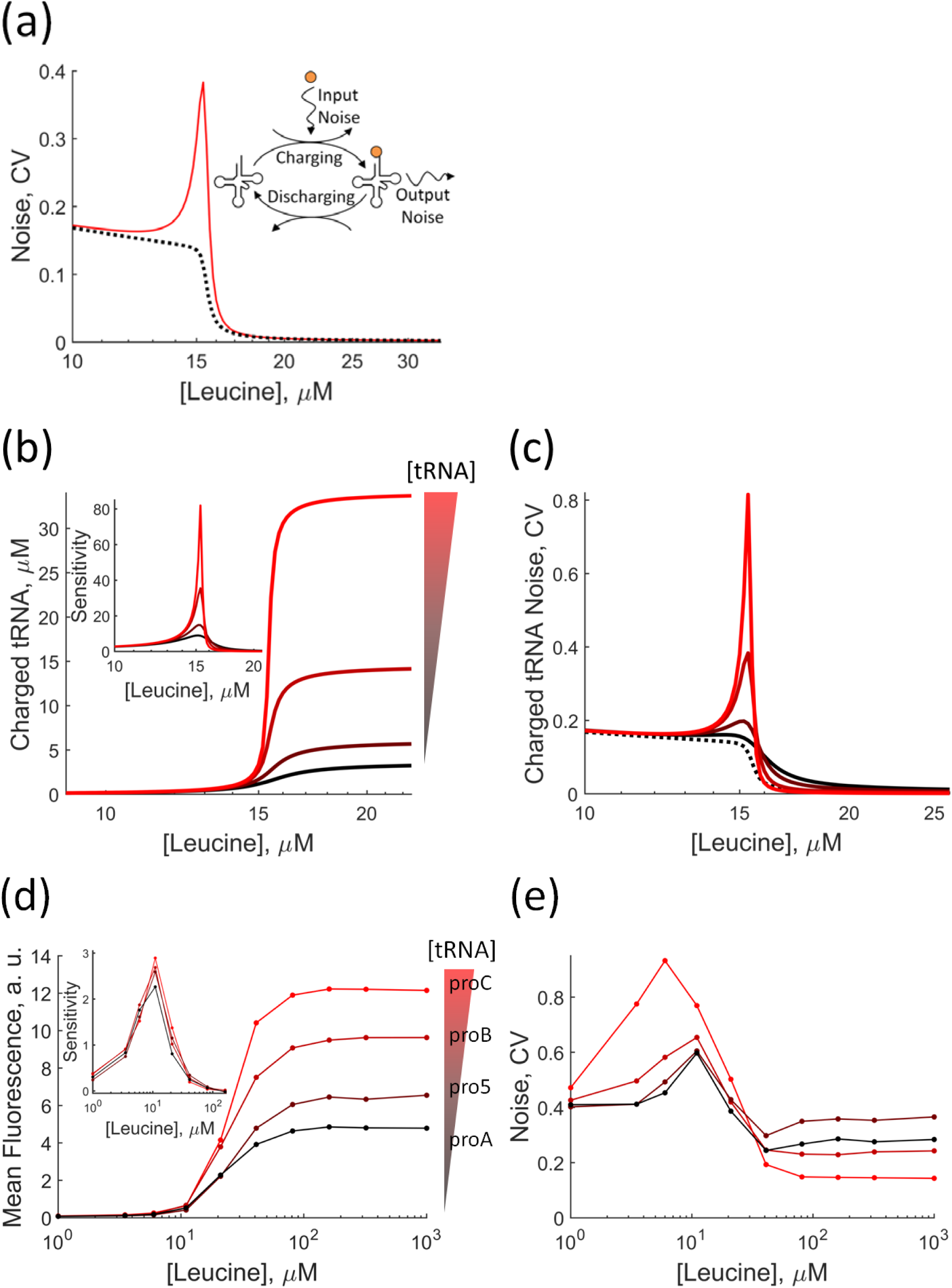
Global fluctuations in the tRNA charging cycle drive codon noise. **(a)** Output noise of the ultra-sensitive charging cycle in the absence **(dashed line)** and the presence **(red line)** of global noise. **Inset**, schematic representation of the hypotheses of the model where global fluctuations, e.g. in synthetase, translating ribosomes or amino acid levels, are amplified through the tRNA charging cycle. **(b)** A minimal charging/discharging tRNA model predicts that, by increasing [tRNA] by 9-fold from minimum (black) to maximum (light red), the maximum fraction of charged tRNA also increases together with amplification sensitivity of the cycle **(Inset). (c, full lines)** Model predictions of the coefficient of variation (noise) associated with the charged fraction of tRNA as a function of [leucine] for different [tRNA] as in **(b)**. **(d)** Mean fluorescence of YFP coded with the sensitive codon CTC as a function of [leucine]. We expressed in Δ*leuU* strains different levels of the essential 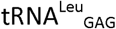, *leuU*, using synthetic promoters with a 9-fold difference in strength from weakest to strongest (black to red). Since CTC is solely read by leuU, [leuU] limits the expression of YFP. **Inset**, evolution of the sensitivity amplification (i.e. logarithmic gain) as a function of available leuU tRNA. **(e)** Noise as a function of leucine concentration for the same strains as in **(d)**. Codon noise was measured as the coefficient of variation of the fluorescence signal from single cells whose means are plotted in **(d)**. Mean and CV values were derived from a sample size of 3×10^3^ ± 200 cells. Error bars associated with the standard error of the mean are smaller than symbols sizes. The model uses previously published rates for the charging and discharging of tRNA, assumes Poisson noise in amino acid fluctuations and uses previously determined parameters for the Michaelis-Menten constants of the synthetase and ribosome activity.

To test our hypotheses and model, we perturbed our system by changing the levels of one specific [tRNA] in order to change the saturation level of the cognate synthetase. Most codons are read by multiple tRNAs, which makes such experiments difficult to design and interpret. We circumvented this challenge by using a YFP construct that uses the sensitive codon CTC that is known to be solely read by 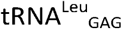 (leuU). This unique tRNA-codon pair makes it possible to use CTC as a readout of leuU charging. To adjust different levels of leuU in the cell and to limit the possibility of compensatory regulation, we deleted *leuU* from the chromosome and expressed it from plasmids under the control of promoters with various strengths spanning a 9-fold range. We hypothesized that as we increase the concentration of tRNAs, the enzyme that catalyzes the amino-acylation works more at saturation, which in turn should increase the amplification sensitivity of the charging/discharging cycle. As predicted, when we increased [leuU] the mean fluorescence increased, which is reflected in the simple model by a similar increase of the charging level (Fig. 4b and 4d). More importantly, we found that the amplification sensitivity (insets in Fig. 4b and 4d) increased with [leuU] and resulted in large expression noise in both the model and experiments (Fig. 4c and 4e).

To better assess the generality of our results, we characterized the noise associated with the isoleucine, threonine and valine codon families. We followed the same experimental strategy as for leucine codons and re-coded the *yfp* reporter using exclusively one specific synonymous codon from the family under study. We built and used auxotrophs for each codon family under study (**Supplementary Note**). We then measured the fluorescence in individual cells after a simultaneous induction of *yfp* expression and a downshift in the cognate amino acid in the media. In all these codon families, we again found that (i) synonymous codons presented varying levels of robustness to amino acid availability and that (ii) the codons most sensitive to starvation produced the highest levels of protein expression noise, while the most robust codons were the least noisy, see **Supplementary Fig. 4**.

In this work, we found that when we shift bacteria from nutrient rich to poor environments, the strength of noise in gene expression can be directly regulated by the choice of synonymous codons used in the genes. The standard model of gene expression noise is based on transcriptional bursts and stochasticity that arises from low numbers of mRNA molecules. By contrast, codon noise is governed by translational processes and does not require low molecular/mRNA numbers (Fig. 1 and 2). Many bacterial processes are “stress-response” programs that are triggered by downshifts in nutrients: such programs include the growth of biofilms, synthesis of flagellum, toxin/anti-toxin systems, and the expression of antibiotic resistance genes. The response to fluctuating growth conditions is also a key challenge for the construction of robust synthetic genetic networks. We hypothesize that this new mode of codon-based noise regulation could be particularly relevant to generating cell-to-cell variability under such nutrient driven adaptive processes.

## Author Contributions

PC conceived and supervised the project. LFM performed preliminary experiments that lead to the effect presented in Fig. 1, 2. EB performed all experiments, computations, designed genetic constructs and wrote the supplementary material. JMK developed the protocol to perform scarless chromosomal deletions. EB and PC wrote the manuscript.

## References

1. J. M. Raser, E. K. O’Shea, Noise in gene expression: origins, consequences, and control. Science 309, 2010–2013 (2005).

2. G. Balazsi, A. van Oudenaarden, J. J. Collins, Cellular decision making and biological noise: from microbes to mammals. Cell 144, 910–925 (2011).

3. J. Paulsson, Summing up the noise in gene networks. Nature 427, 415–418 (2004).

4. E. M. Ozbudak, M. Thattai, I. Kurtser, A. D. Grossman, A. van Oudenaarden, Regulation of noise in the expression of a single gene. Nat Genet 31, 69–73 (2002).

5. I. Golding, J. Paulsson, S. M. Zawilski, E. C. Cox, Real-time kinetics of gene activity in individual bacteria. Cell 123, 1025–1036 (2005).

6. A. R. Subramaniam, T. Pan, P. Cluzel, Environmental perturbations lift the degeneracy of the genetic code to regulate protein levels in bacteria. Proc Natl Acad Sci U S A 110, 2419–2424 (2013).

7. J. Elf, D. Nilsson, T. Tenson, M. Ehrenberg, Selective charging of tRNA isoacceptors explains patterns of codon usage. Science 300, 1718–1722 (2003).

8. M. B. Elowitz, A. J. Levine, E. D. Siggia, P. S. Swain, Stochastic gene expression in a single cell. Science 297, 1183–1186 (2002).

9. A. Hilfinger, J. Paulsson, Separating intrinsic from extrinsic fluctuations in dynamic biological systems. Proc Natl Acad Sci U S A 108, 12167–12172 (2011).

10. G. Chalancon et al., Interplay between gene expression noise and regulatory network architecture. Trends Genet 28, 221–232 (2012).

11. J. Elf, M. Ehrenberg, Fast evaluation of fluctuations in biochemical networks with the linear noise approximation. Genome Res 13, 2475–2484 (2003).

12. O. G. Berg, J. Paulsson, M. Ehrenberg, Fluctuations and quality of control in biological cells: zero-order ultrasensitivity reinvestigated. Biophys J 79, 1228–1236 (2000).

13. J. Elf, J. Paulsson, O. G. Berg, M. Ehrenberg, Near-critical phenomena in intracellular metabolite pools. Biophys J 84, 154–170 (2003).

14. J. Elf, M. Ehrenberg, Near-critical behavior of aminoacyl-tRNA pools in E. coli at rate-limiting supply of amino acids. Biophys J 88, 132–146 (2005).

15. J. Elf, O. G. Berg, M. Ehrenberg, Comparison of repressor and transcriptional attenuator systems for control of amino acid biosynthetic operons. J Mol Biol 313, 941–954 (2001).

