## Supplementary material for "Codon-dependent noise dictates cell-to-cell variability in nutrient poor environments"

Enrique Balleza et al,.

### 1 Construction of Auxotrophs and YFP Synonymous Codon Variants

#### 1.1 General Chromosome Editing Protocols and Construction of the Background Strain, *Noise bg V1.0*.

We used MG1655 (Coli Genetic Stock Center (CGSC) #6300) as our background strain. We constructed all derived strains following standard lambda red recombination protocols using the pSIM5 helper plasmid^1^. When needed, we performed scarless chromosomal engineering^2^ using the selection/counterselection cassette kan-ccdB in pKD45^3^. First, we inserted the kan-ccdB cassette at the desired locus. Then, using counterselection against the ccdB marker, we inserted the final desired construct. We also used scarless chromosomal engineering to avoid inserting into the chromosome unnecessary antibiotic resistance cassettes.

We inserted a cassette to constitutively express a cyan fluorescent protein (CFP) at position 84.3 min of our background strain^4^. The fluorophore served as a marker to separate cells from debris in flow cytometry experiments. The background strain also contained a cassette with a copy of the tetR protein and the SpecR antibiotic resistance gene at the AttB site. The cassette was derived from plasmid pZS4^5^. We designated this strain Noise bg V1.0. To insert the different recoded versions of the YFP reporter (YFP starvation reporters), we used the SpecR as a *landing pad*: we substituted the SpecR gene with a cassette containing the desired recoded YFP-AmpR cassette.

#### 1.2 Construction of YFP Synonymous Codon Variants.

The promoter of the reporter is P_LtetO_^5^. This promoter is strongly repressed in strain Noise bg V1.0 by the tetR protein and can be induced by addition of anhydrotetracycline (aTc). Translation initiation is controlled by Variant#1, a strong ribosomal binding site (RBS) derived from gene 10 of bacteriophage T7^6^. Given that sequence recoding near the RBS region affects translation initiation rate by the strengthening/relaxation of mRNA hairpins^7,8^, we did not recode with synonymous codons the first 78 nucleotides after the start codon of our YFP reporter. This strategy guaranties that differences in protein synthesis are due to codon substitutions rather than to a change in the affinity of the 5’end coding sequence for the RBS. In the YFP reporter, the number of recoded synonymous codons for each different amino acid family is: isoleucine, 9 codons; leucine, 6 codons; threonine, 10 codons; and valine, 9 codons. All sequences are available as Supplementary Material.

#### 1.3 Auxotroph Construction

Depending on the desired auxotrophy, we deleted from strain Noise bg V1.0 an essential amino acid biosynthesis enzyme. We used ccdB counterselection to avoid a remnant antibiotic cassette after the deletion. The final strains contained a scar with the first and last 39 nucleotides from the deleted gene except for the leucine auxotroph, ΔleuB, for which the scar corresponds to the chromosomal homology of the Keio collection^9^ knockout primers.

In this work, we used the following amino acid auxotrophs (gene deletions): valine (Δ*ilvC*), threonine (Δ*thrC*), isoleucine (Δ*ilvE*), leucine (Δ*leuB*).

### 2 Test of Auxotrophy and Valine, Threonine, Isoleucine & Leucine Downshift Experiments

#### 2.1 Media Composition

We prepared amino acid dropped-out solutions using Type II water (ASTM Type II water RO/DI, EMD 7732-18-5). We purchased all amino acids from Sigma. The source of amino acids was non-animal except for L-asparagine (Sigma A4159-25G), L-aspartic acid (Sigma A4534), L-tyrosine (Sigma 93829), and L-lysine (Sigma 62929). We prepared 10X concentrates of dropped-out solutions and adjusted the pH (7-8) with NaOH. We filter-sterilized, aliquoted and froze (-20°C) freshly prepared stocks. The milimolarity of the different L-amino acids in a 1X solution was: alanine (0.5), arginine (0.2), asparagine (0.3), aspartic acid (0.3), cysteine (0.4), glutamic acid (0.3), glutamine (0.3), glycine (0.6), histidine (0.2), isoleucine (0.3), leucine (0.3), lysine (0.2), methionine (0.3), phenylalanine (0.3), proline (0.4), serine (0.4), threonine (0.4), tryptophan (0.2), tyrosine (0.2), valine (0.4).

The 1X composition of our base media was: MOPS (Teknova M2101), 1.320 mM K2HPO4 (Teknova M2102), 0.2% Glucose, 1X dropped-out amino acid solution. Prior to starvation of auxotrophs, we supplemented, with the corresponding dropped-out amino acid, the media for overnights and the media for exponential growth using the following final concentrations: 1mM L-leucine, 1mM L-threonine, 1mM L-isoleucine, 1mM L-valine.

#### 2.2 Auxotrophy Test

To test auxotrophy, we set cultures of auxotrophs in drop-out media (MOPS, 1.320 mM K2HPO4, 0.2% Glucose, 1X drop-out amino acid solution; see also subsection **Media Composition**) without the corresponding amino acid and incubated them at 30°C, 250rpm shaking. The cultures were set by making a 1:1500 dilution from a single colony resuspension in LB. We also incubated a positive control (media supplemented with cognate amino acid and corresponding auxotroph) and a negative control (media without cells and without cognate aminoacid). We performed the auxotrophy test twice. After 24 hrs of incubation, all different auxotrophs did not grow in media lacking the cognate amino acid. For ΔserA, we followed the same auxotrophy test except that we also dropped out glycine from the media, see **Supplementary Figure 1**.

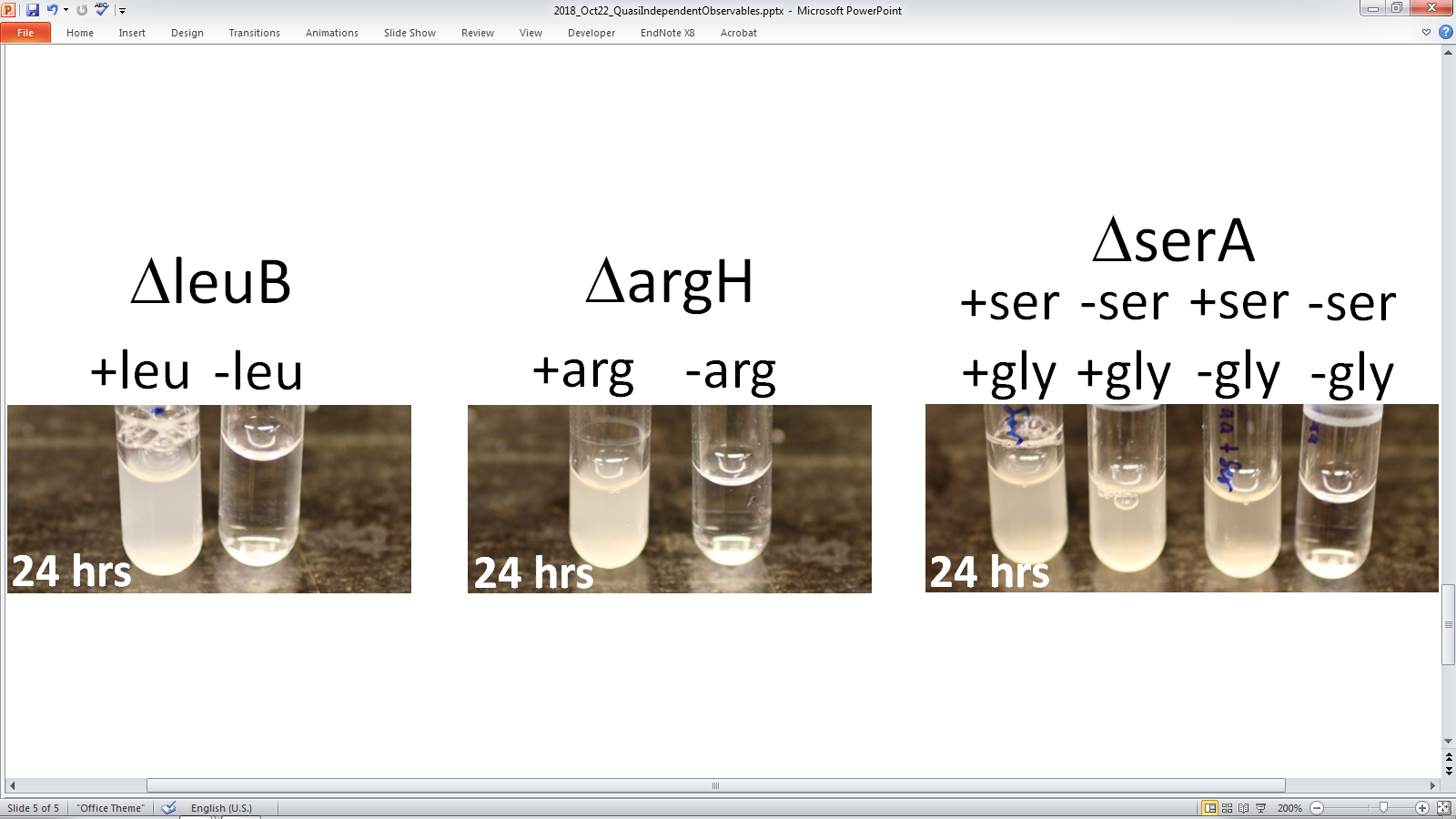

**Supplementary Figure 1** | **Auxotrophy test example.** An MG1655 strain derivative with a deletion of the *leuB* gene grown in dropout media with and without leucine (1mM). After 24 hrs of growth at 30°C and 250 rpm shaking, only the culture with leucine grows. See subsection **Media Composition** for media composition details.

#### 2.3 Protocol to Estimate Basal Translation Activity during Starvation

We aliquoted 1ml of “starvation media” (MOPS media supplemented with the amino acid dropout mixture of interest, see subsection **Media Composition**) into twelve wells of a 2 ml 96-well plate. We also prepared a 20X solution of aTc (4ug/ml) in ASTM Type II water and aliquoted 55ul into twelve wells of a microtiter 96-well plate. We started a fresh culture doing a 1000X dilution from an overnight culture; both cultures were grown in “starvation media” supplemented with all amino acids. After 4.5-5 hrs of growth (250rpm, 30°C), we transferred 1ul of culture into 1ml of the “starvation media” using a 12-channel pipette in order to do a simultaneous transfer into all the twelve wells of the 2 ml 96-well plate. Immediately, we added 50ul of the 20X aTc solution into the first well and incubated the 2 ml 96-well plate (1300 rpm, 30°C). After 15 min, we transferred 240ul of sample from the 1st well into a well of a microtiter 96-well plate and stored the plate at 4°C. We performed the transfer using prechilled pipette tips and a prechilled microtiter 96-well plate. This first transfer would correspond to time “zero” of the amino acid down shift. We similarly interrogated other time points with the remaining starving parallel cultures, accumulating samples in the same prechilled 96-well plate. After 3 to 4hrs of saving samples in the same prechilled 96-well plate, we measured fluorescence using the protocol described in the **Flow Cytometry** section. For the samples stored in the prechilled 96-well plate, we did not observe any dependence in fluorescence when we measured fluorescence within a range of 1 to 6hrs.

#### 2.4 Basal Translation Activity after an Amino Acid Downshift

We devised a protocol to quantify the translation activity after an amino acid down shift, see **Protocol to Estimate Basal Translation Activity during Starvation**. In brief: We inoculated drop-out media with cells previously growing exponentially in amino acid rich media. We performed the inoculation in 12 different wells of a 2ml deep, 96-well plate using a multichannel pipette. In this way, we simultaneously performed an amino acid downshift in many parallel cultures. To interrogate at a specific time point the translation basal activity, we sacrificially induced a culture with aTc for 15 min. Then, we transferred an aliquot into a pre-chilled test tube to later estimate population size and translation activity using flow cytometry, see section **Flow Cytometry** for details. We repeat this process with cultures induced at different time points.

As a reference experiment, we quantified the autofluorescence of our leucine auxotroph strain without the P_LtetO_-YFP cassette. Also, as a control experiment, we measured the repression level of promoter P_LtetO_ in the context of constitutive TetR expression and no aTc. We performed this control using our leucine auxotroph strain bearing the P_LtetO_-YFP_CTG_ cassette. For the autofluorescence case, we noticed a very low and constant level of mean fluorescence throughout several hours. This was also the case for the repression strength control and, as expected, the mean fluorescence was marginally higher than in the autofluorescence control with the strain that had no P_LtetO_-YFP cassette (**Supplementary Figure 2**, 1^st^ row, left column).

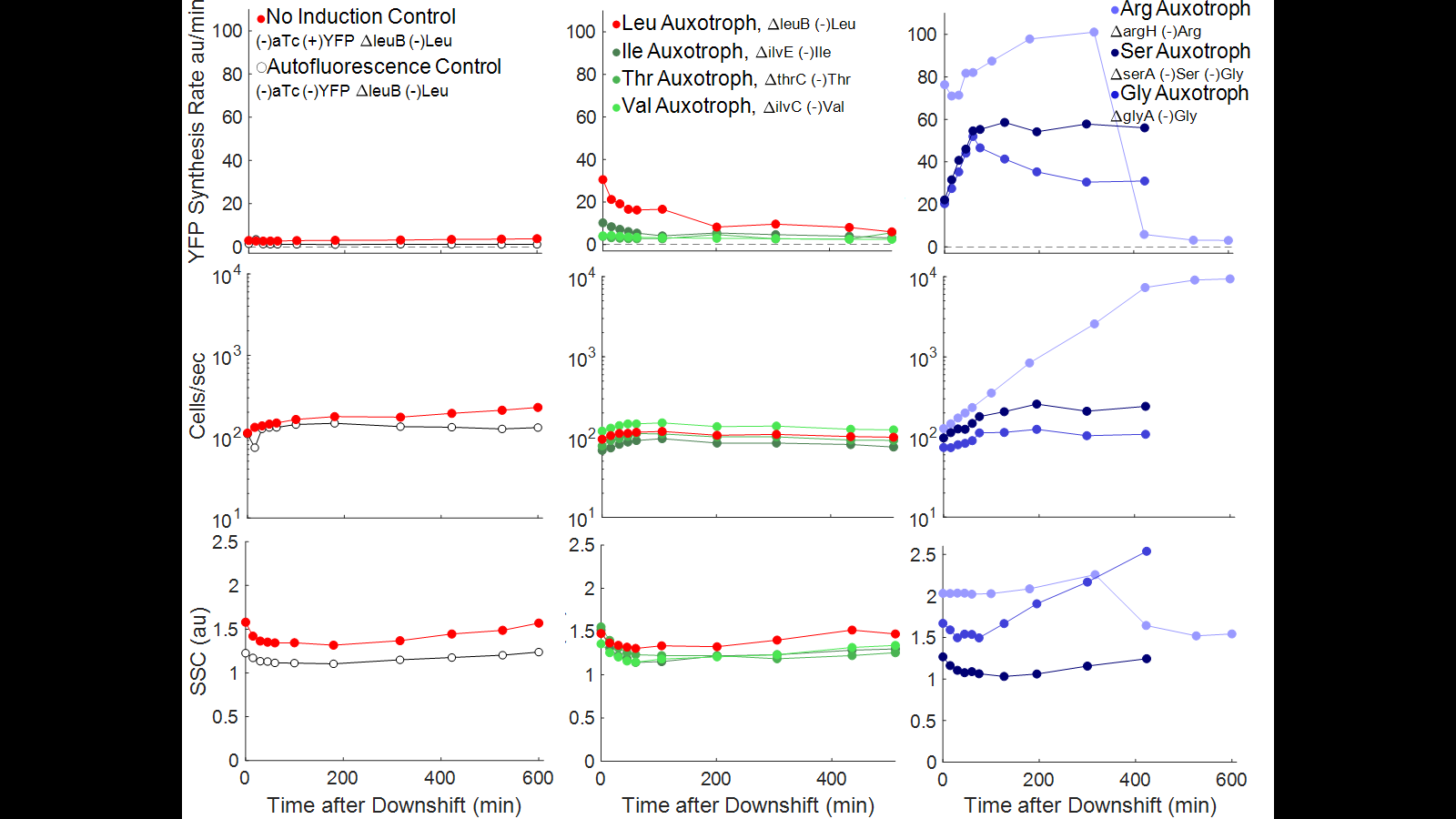

**Supplementary Figure 2** | **Evolution of basal translation activity, number of cells and side scattering during amino acid downshift for seven different auxotrophs.** **Top row.** Mean YFP synthesis rate estimated by sacrificially inducing, for 15 min, a sample culture at different time points after amino acid downshift. **Middle row.** Cells per second as a proxy for cell division rate: To obtain the number of cells per second, we held constant for all data points the sample flow rate in the High Throughput Sampler of the flow cytometer then, for each data point, we divided the number of detected cells by the time interval defined by the first and last cells detected. **Bottom row.** Average side scattering as an indicator of change in cell geometry. **Left column.** Open circles correspond to an autofluorescence control (Noise bg V1.0, no YFP, no aTc added). Red circles correspond to a control to assess strength of repression (Leu auxotroph, YFP_CTG_, no aTc added). **Central column.** Leu auxotroph (Δ*leuB*, YFP_CTG_, no leucine) in red circles; Ile auxotroph (Δ*ilvE*, YFP_ATC_, no isoleucine) in dark green circles; Thr auxotroph (Δ*thrC*, YFP_ACC_, no threonine) in green circles; and Val auxotroph (Δ*ilvC*, YFP_GTG_, no valine) in light green circles.

We quantified for both strains, with and without P_LtetO_-YFP cassette, the cell population evolution throughout the duration of the experiment. As expected, the auxotrophs did not divide nor lysed (**Supplementary Figure 2**, 2^nd^ row, left column). This result is along the same line as that by Flint^10^ who recorded *E coli*‘s starvation-survival in autoclaved river water over periods of time up to 260 days, and who also found that the number of *E. coli* cells remained constant.

As a final quantification, we used the mean of the side scattering (SSC) as an indicator of changes in cell shape. It has been well documented that forward and side scattering can be used to reliably estimate the size of submicron spherical particles (see article^11^ by Poncelet *et al.* for the discrimination of different populations of submicron spheres with the Fortessa flow cytometer, the model used in this study). However, the shape of *E. coli* is cylindrical and can range from spherical to filamentous, which makes a precise size estimation difficult using SSC. Thus, we restrict ourselves to use changes in the mean SSC as a qualitative indicator of changes in cell shape without further quantification. In the autofluorescence and repression strength controls, we inferred that there was basically no change in cell shape because the SSC mean value remained constant (**Supplementary Figure 2**, 3^rd^ row, left column). The cell fluorescence, population and shape evolution results, taken together, indicate that the control strains are neither dividing, growing nor lysing, and that the amount of auto-fluorescence and leaky expression is negligible.

The second column of **Supplementary Figure 2** shows the basal translation activity, of the threonine and of the branched-chain amino acids (BCAA) auxotrophs—leucine, isoleucine and valine. Again we can see that these auxotrophic strains do not divide nor grow. The translation basal activity is more interesting and supports the hypothesis that, as a first approximation, the internal concentration of threonine and BCAAs after a downshift is minimal. An equivalent hypothesis is that, when cells are growing exponentially and before the downshift, the influx and the consumption of amino acids is balanced. For leucine, after the downshift, a closer examination shows that there is a small concentration of internal amino acid that decays in about an hour. In general, results in the second column of **Supplementary Figure 2** suggest that the threonine and BCAAs auxotrophs respond immediately to a change in external amino acid concentration, see section **Starvation Experiments of Other Codon Families**.

#### 2.5 Downshift Protocol and Downshift Experiments

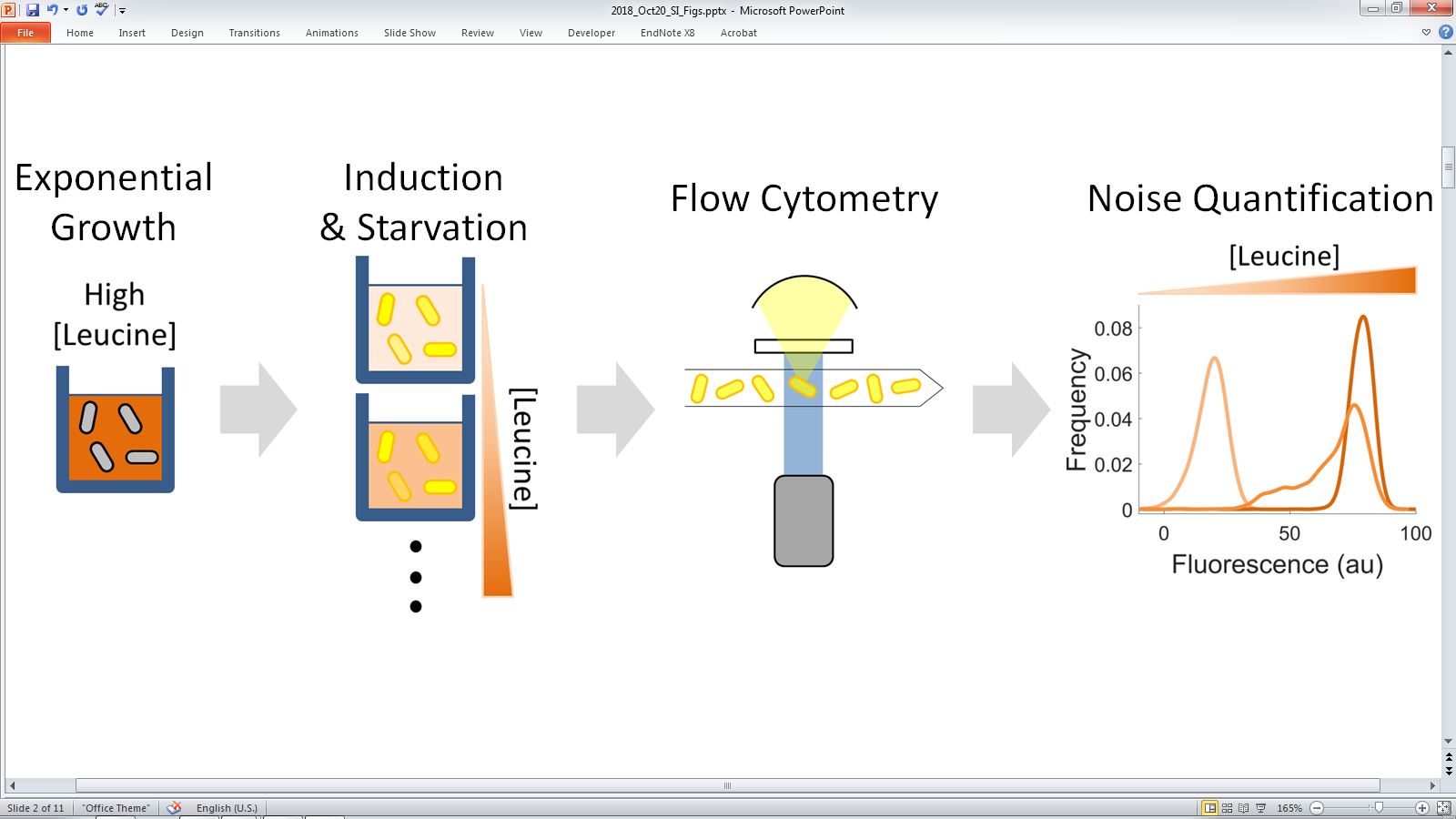

**Supplementary Figure 3** | **Schematic of the amino acid downshift protocol.** From left to right. (i) We grew a culture until early exponential phase of an auxotroph in media supplemented with the cognate amino acid. (ii) After 4.5-5hrs of exponential growth, we transferred 1ul of culture into 1ml of media supplemented with inducer and different concentrations of the cognate amino acid and incubated for 1hr. (iii) Then, for the different amino acid concentrations, we quantified the fluorescence from the induced YFP synonymous codon variant using a flow cytometer. (iv) Finally, we automatically separated cells from debris using a CFP marker, obtained the fluorescence distribution of the samples and calculated the fluorescence mean and coefficient of variation.

The protocol to obtain the response to a downshift in isoleucine, leucine, threonine and valine was the following: The night before the experiment, we grew the auxotroph of choice carrying the YFP cassette with synonymous substitutions of the cognate codon in our base media supplemented with its cognate amino acid (1mM), see subsection **Media Composition**. The day of the experiment, we performed a 1000X dilution of the overnight into 3ml of our base media supplemented with 1mM of cognate amino acid. We grew the auxotroph for 4.5-5hr at 30°C, 250 rpm. Then, we transferred 1ul of culture into 1ml of base media with a predefined concentration of cognate amino acid and inducer (aTc, 0.2ug/ml) and incubated for 1hr at 30°C with shaking at 1350 rpm (see below). Finally, we transferred 240ul of culture into pre-chilled 96 microtiter wells and measured single-cell fluorescence using a flow cytometer, see section **Flow Cytometry**.

In order to start simultaneously all cultures with different amino acid concentrations, we poured the 3ml culture into a reagent reservoir (VWR, 89094-662) and, using a 12-channel pipette, we transferred 1ul of culture into 11 wells of a 2ml deep, 96-well plate. About 1hr before the simultaneous transfer, we prepared the 2ml deep, 96-well plate with predefined amino acid concentrations. The transfer procedure was performed at 30°C and the plate incubated for 1hr at 30°C and shaken at 1350 rpm using a Titramax 100 (Heidolph Instruments). Prior to the transfer procedure, the pipette tips and the reagent reservoir, as well as the 2ml, 96-well plate with media were thermalized to 30°C.

The micromolarity of the tested amino acid concentrations were for isoleucine 1+(0, 2.5, 5, 10, 15, 20, 40, 80, 160, 360, 1000); for leucine 1+(0, 2.5, 5, 10, 15, 20, 40, 80, 160, 360, 1000); for threonine 1+(0, 5, 10, 20, 40, 80, 160, 320, 1000, 2000, 4000); and for valine 1+(0, 5, 10, 20, 40, 80, 160, 320, 1000, 2000, 4000). Note that for isoleucine, leucine, threonine and valine ranges we added 1 micromolar to account for the approximate amount of amino acid transferred from the 1000X dilution of the exponential culture which had [cognate amino acid] = 1mM. The auxotrophic strains were: for valine, ΔilvC; for threonine, ΔthrC; for isoleucine, ΔilvE; for leucine, ΔleuB.

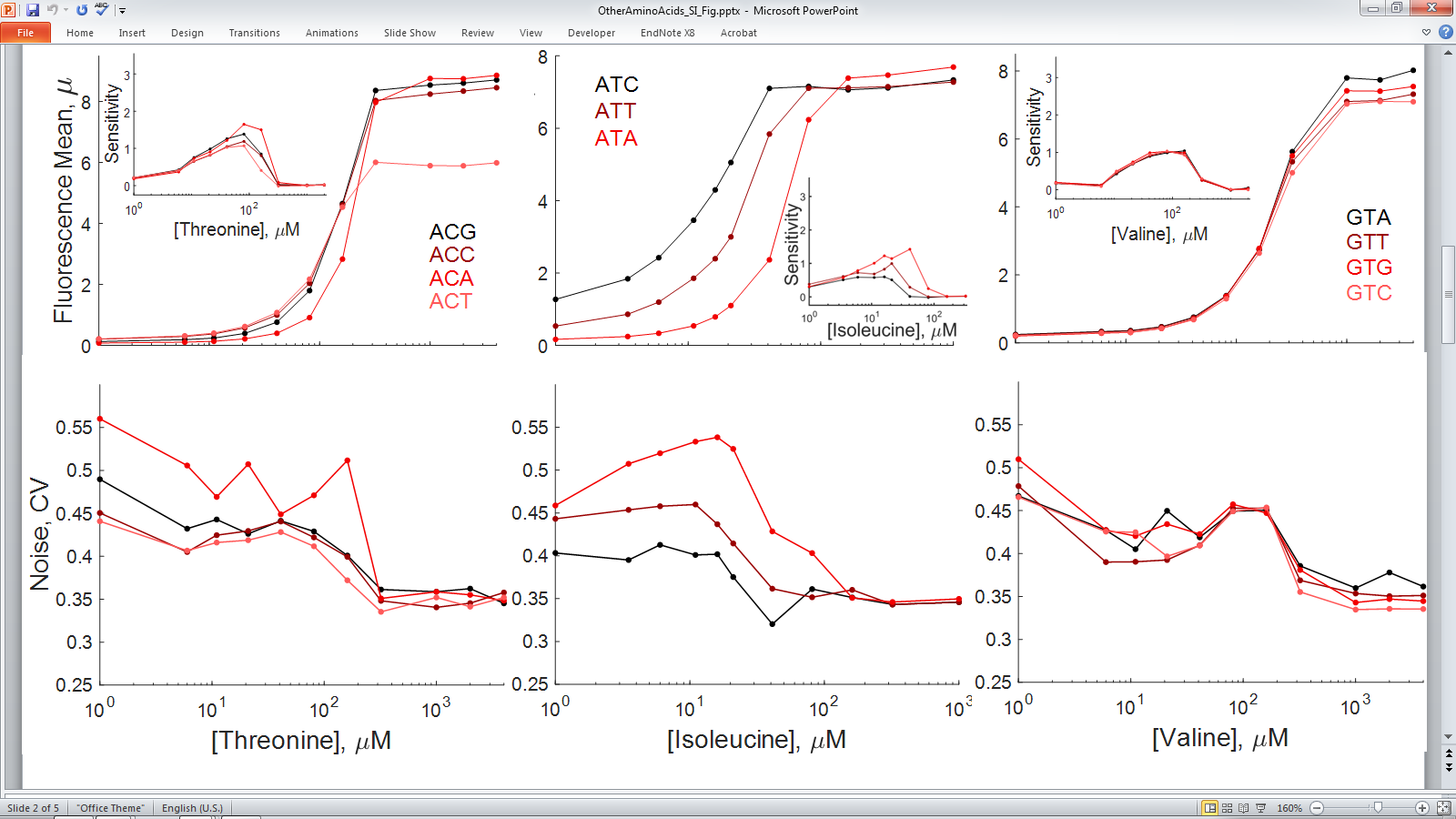

**Supplementary Figure 4** | **The choice of synonymous codons from different codon families underlies translation noise during cognate amino acid limitation.** Mean fluorescence (1^st^ row) and associated noise (2^nd^ row) of a YFP starvation reporter as a function of the cognate amino acid concentration, and the synonymous codon choice of the YFP reporter for, from left to right, the threonine, isoleucine and valine codon families. Insets show the sensitivity amplification or logarithmic gain of the mean fluorescence curves, $(\Delta\mu/\mu)/(\Delta AminoAcid/AminoAcid)$. Note how high sensitivity amplification is associated to large random fluctuations (high CV) and how for every codon the sensitivity amplification correlates with the noise level of the codon.

#### 2.6 Leucine Replicate Experiment

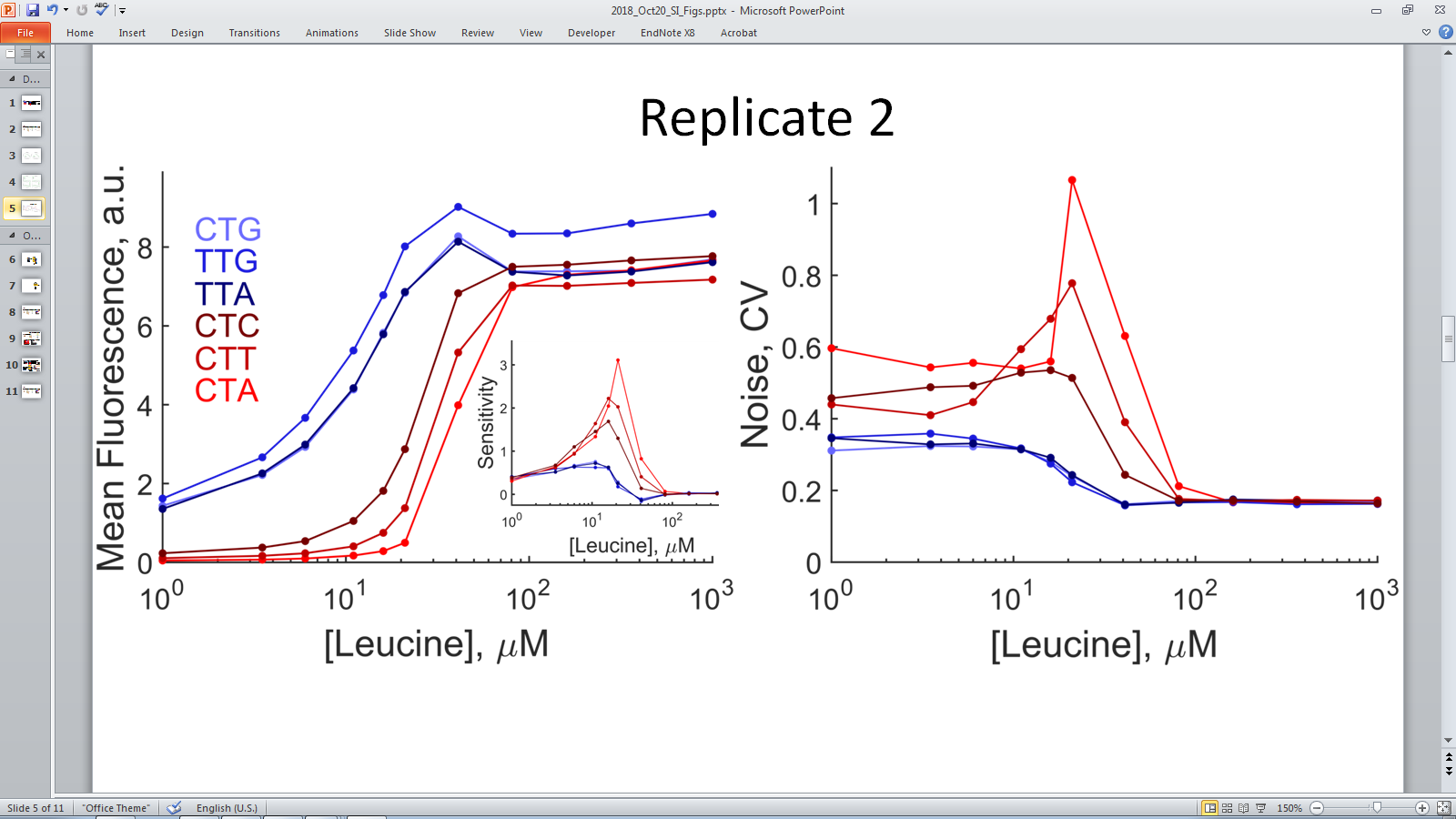

**Supplementary Figure 5** | **Replicate of figure 2 of main text.** Mean fluorescence (left) and associated noise (right) of a YFP reporter as a function of the concentration of leucine and the synonymous codon choice of the YFP reporter for the leucine codon family. The inset shows the sensitivity amplification or logarithmic gain of the mean fluorescence curves, $(\Delta\mu/\mu)/(\Delta Leu/Leu)$. Mean and CV values were derived from a sample size of 3x10^3^ ± 200 cells. Error bars associated with the standard error of the mean are smaller than symbols sizes. The data from this experiment corresponds to a different set of replicate strains of the YFP variants of synonymous codons of the leucine family.

### 3 Flow Cytometry

#### 3.1 Data Acquisition

We sampled cultures from microtiter 96-well plates with a Becton Dickinson High Throughput Sampler (BD HTS) using the standard mode and the following parameters: Sample flow rate 0.5 uL/sec, sample volume 145 uL, mixing volume 50 uL, mixing speed 70 uL/sec, number of mixes 2 cycles, wash volume 800 uL. We performed cytometry using a BD LSR Fortessa. The excitation/emission configuration was CFP Ex 440 (laser), Em 470/20; YFP Ex 488 (laser), Em 542/27. In order to wash the nozzle of the BD HTS and avoid contamination arising from the carryover of highly induced cell cultures into lowly induced cultures, we filled an in-between well with media and without cells; e.g. to minimize the carryover from the highly induced cultures ([leucine]=1mM) in the 96-well plate position (column=11, row=n) into the lowly induced culture ([leucine]=1uM) in well (column=1, row=n+1), we filled well (column=12, row=n) with media and without cells. Occasionally, there were still few highly induced cells contaminating column=1 or 2. In this case, we gated (see subsection **Data Analysis** for a general description on the gates applied to data) only column=1 and 2 with the threshold YFP = 5000au. This threshold is about one order of magnitude above the average expression level at low amino acid concentrations for sensitive codons.

For two-color experiments, we performed cytometry using a BD FACS Aria.

#### 3.2 Data Analysis

We performed the statistical analyses of noise quantification using directly the raw data from the flow cytometry files. For strains expressing constitutively a cyan or a red fluorescent protein (CFP or RFP) as a marker, we gated data using the CFP or the RFP channel and the Side Scatter channel (SSC). The constitutive expression of an FP allowed us to clearly separate debris from cells and to gate consistently data using the same thresholds across different amino acid concentrations or different starvation times. Occasionally, we gated the YFP channel corresponding to the two lowest amino acid concentrations with YFP<5000au to exclude one or two highly induced cells that were carried over, during data acquisition, from conditions of high [amino acid] into conditions of low [amino acid]. For every experiment, all thresholding parameters have been clearly defined in the corresponding Matlab scripts used to generate the main and supplementary figures.

Using the law of total variance, we conditioned on SSC to circumvent the effect of cell size on YFP noise quantification^12,13^,

$$Var\left( YFP \right)=\underset{\begin{aligned} Variation exclusively \\ due to YFP expression \end{aligned}}{\underbrace{\left\langle Var(YFP|SSC) \right\rangle}}\underset{\begin{aligned} Variation of YFP \\ due to SSC \end{aligned}}{+\underbrace{Var(\left\langle YFP|SSC \right\rangle)}}$$

Thus, when we report the YFP coefficient of variation (CV), what we report is the estimated CV exclusively due to noise in YFP expression, i.e. $CV= \left\langle Var\left( YFP | SSC \right) \right\rangle/\langle YFP\rangle$ . We also noted that $Var(\left\langle YFP|SSC \right\rangle)$ contributes 0.2-0.3 CV units to $Var\left( YFP \right)$ and that its value is basically independent of the amount of amino acid concentration (leucine).

In $\left\langle Var(YFP|SSC) \right\rangle$, the conditioning on SSC is analogous to the use of FP concentration in microscopy images of *E. coli* in the sense that both approaches try to eliminate noise contributions from the cell-to-cell variations in DNA content by using, as a proxy for DNA content, cell size; , *i.e.* total fluorescence is on average greater in lager cells , which in turn have on average more DNA content (chromosomes or plasmids).

All data was analyzed with custom Matlab (R2017b) code with the following workflow: reading data from raw files, gating, calculation of statistics, and plotting. Only for data visualization purposes, we used the Vlog transformation^14^ in **Fig. 3a**. All software is available as Supplementary Material.

### 4 Comparison of Codon Noise to Transcription and Translation Initiation Noise

First, we describe the model of constitutive, unregulated gene expression. Then, we derive the expected behavior of the CV when the translation initiation rate changes or when the transcription rate changes and compare with experimental results.

#### 4.1 Model of Gene Expression from an Unregulated Promoter

Following Paulsson^15^, the gene expression model is

| Transcription: $x_{1}\underset{\to}{\lambda_{1}}x_{1}+1$ Translation: $x_{2}\underset{\to}{\lambda_{2}x_{1}}x_{2}+1$  mRNA degradation: $x_{1}\underset{\to}{x_{1}/\tau_{1}}x_{1}-1$ Proteolysis: $x_{2}\underset{\to}{x_{2}/\tau_{2}}x_{2}-1$ | (1) |
| --- | --- |

Where $x_{1}$and $x_{2}$ represent the number of mRNAs and green fluorescent proteins (GFP) respectively, $\lambda_{1}$is the synthesis rate of mRNAs, $\lambda_{2}$ is translation initiation rate of GFP per mRNA, and $\tau_{1}$ and $\tau_{2}$ are the average lifetimes of mRNAs and GFPs respectively. Using the linear noise approximation^15,16^, the stationary and normalized fluctuations in the number of proteins is

$$\frac{\sigma_{2}^{2}}{\left\langle x_{2} \right\rangle^{2}}=\frac{1}{\langle x_{2}\rangle}+\frac{1}{\left\langle x_{1} \right\rangle}\frac{\tau_{1}}{\tau_{1}+\tau_{2}}$$

(2)

For the model we are considering, this solution is exact because all rates are constant or linear. Given that GFP is a stable protein, $\tau_{1}\ll\tau_{2}$, the previous equation reduces to

$$CV^{2}=\frac{\sigma_{2}^{2}}{\left\langle x_{2} \right\rangle^{2}}=\frac{1}{\langle x_{2}\rangle}+\frac{1}{\left\langle x_{1} \right\rangle}\frac{\tau_{1}}{\tau_{2}}$$

(3)

Where $\sigma_{2}^{2}$ is the protein variance, and $CV$is the coefficient of variation.

#### 4.2 Comparison of Codon Noise to Transcription Noise

Using the macroscopic rate equations, $d\langle x_{1}\rangle/dt=\lambda_{1}-\langle x_{1}\rangle/\tau_{1}$ and $d\langle x_{2}\rangle/dt=\lambda_{2}\langle x_{1}\rangle-\langle x_{2}\rangle/\tau_{2}$, and assuming stationarity, we can write the last equation as

$$CV^{2}=\frac{1}{\left\langle x_{2} \right\rangle}(1+\lambda_{2}\tau_{1})$$

(4)

In terms of the average lifetimes and rates, the average protein number, $\left\langle x_{2} \right\rangle=\lambda_{1}\tau_{1}\lambda_{2}\tau_{2}$ , depends linearly on the transcription rate, $\lambda_{1}$. Thus, the $CV^{2}$ is proportional to ${1/\lambda}_{1}$, or $CV\propto\left\langle x_{2} \right\rangle^{-1/2}$. This relationship has been shown to hold between the expression of different proteins and the associated noise under different experimental conditions^17,18^. Here, we are interested in testing if equation (4) holds in *E. coli* under different concentrations of a limiting amino acid (leucine). Afterwards, we compare the magnitude of transcription noise to the magnitude of codon noise.

We compared relationship (4) with flow cytometry data from a chromosomal copy of YFP (a leucine starvation reporter encoded with a robust CTG codon) under the control of P_LtetO_, a Tet inducible promoter. To modulate the transcription rate, we varied the aTc concentration. We observed that the relationship, $CV\propto\left\langle x_{2} \right\rangle^{-1/2}$, held for about one order of magnitude in mean fluorescence levels, see **Supplementary** **Figure 6**.

**Strains, experimental protocol and replicates.** We estimated transcription noise using the YFP variant encoded with the robust codon “CTG” (a member of the set of the leucine codon YFP variants). The gene of this variant was inserted at the AttB site of our leucine auxotroph background strain (Noise bg V1.0, ΔleuB), see **Construction of Auxotrophs and YFP Synonymous Codon Variants**. To modulate transcription rate, we modified the amino acid downshift protocol in **Section 2.5**.The goal of this protocol was to quantify transcription noise for different aTc concentrations at a fixed level of starvation. First, we prepared rows of a 2ml deep, 96-well plate with starvation media with different leucine concentrations; [leucine] = 10, 15, 20, 40 and 80uM in each row. Additionally, we filled wells in a row of a microtiter 96-well plate with 55ul of aTc with concentrations that started at a 20X aTc concentrate (4ug/ml) and were followed by seven sequential two-fold dilutions; we prepared in parallel as many of these rows with “aTc sequential dilutions” as the number of different tested leucine concentrations. Then, we transfer a row of “aTc sequential dilution” to a row of starvation media. We repeated this transfer for all different amino acid concentrations. From this point onwards, we followed the incubation of cells in starvation media as described in **Section 2.5**. We repeated the response to different [leucine] and different [aTc] twice using the same YFP robust CTG codon variant strain, see **Supplementary** **Figure 6**.

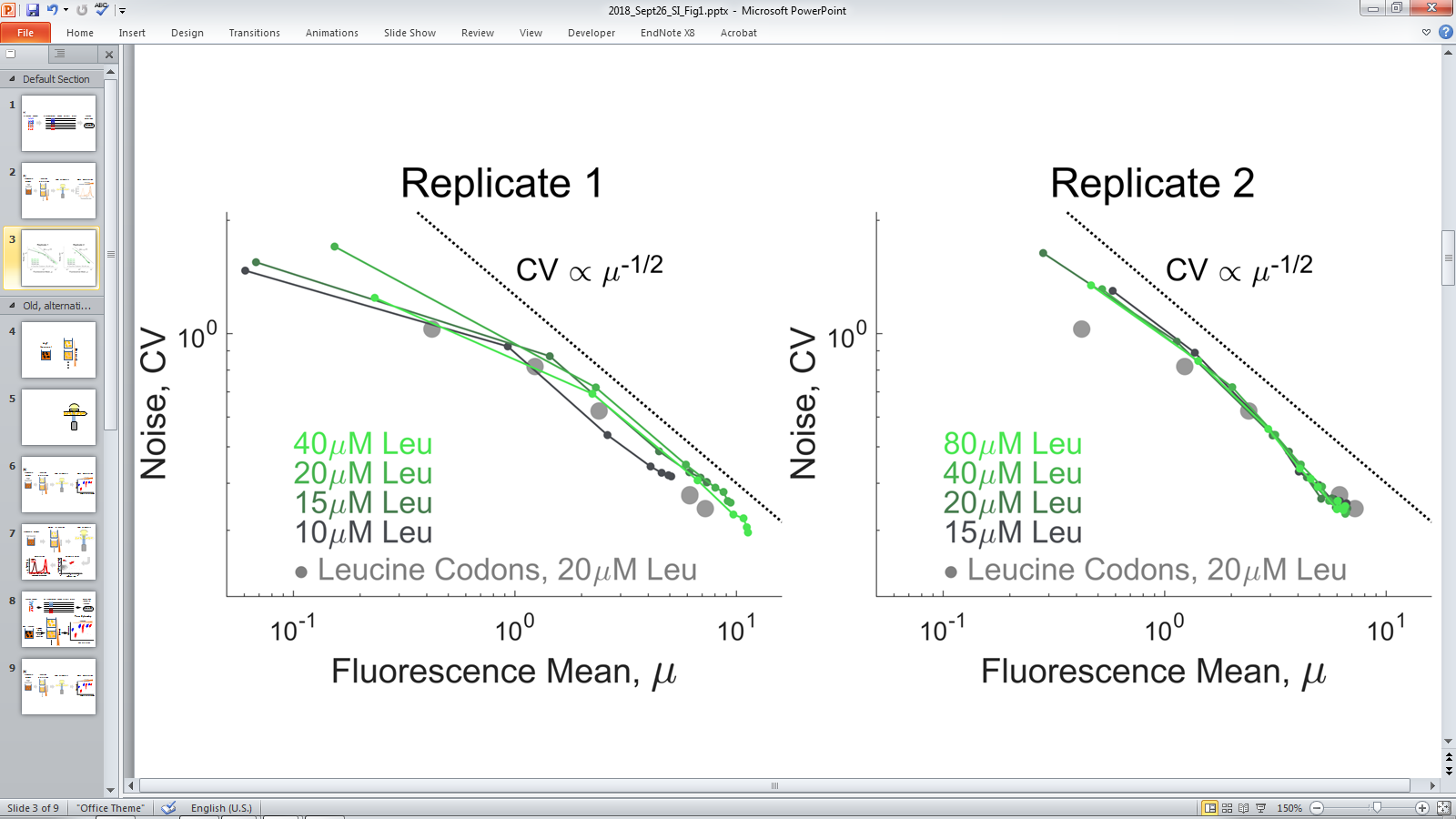

**Supplementary** **Figure 6** | **Experimental relationship between mean and associated noise of the CTG starvation reporter at different transcription rates—*i.e.* different aTc levels—and different fixed leucine concentrations.** Shades of green indicate different leucine concentrations. To modulate transcription rate, we incubated cells expressing the starvation reporter CTG at different concentrations of the aTc inducer covering a ~70X induction range. In each starvation conditions, the highest aTc concentration corresponds to the rightmost data point. We chose different leucine concentrations within the range at which leucine codon noise displays a maximum, see **Fig. 2**. Note that the range of [leucine] is different between the two repeats. Data in gray circles is the noise associated to leucine synonymous codons when codon noise is maximal at [leucine]=20uM, see **Fig.2**. Codon noise data in gray circles is also the same as that presented in **Fig. 1**. The dashed line indicates the expected relationship between mean protein and associated noise.

#### 4.3 Comparison of Codon Noise to Translation Noise

Given what it is known about the protein-to-mRNA ratio in *E. coli* cells—an average mRNA produces 1,000 proteins per division cycle^19^–we can assume that, for the typical *E. coli* protein, $x_{2}$, the first noise term in eq. (3),$CV^{2}=\frac{1}{\langle x_{2}\rangle}+\frac{1}{\left\langle x_{1} \right\rangle}\frac{\tau_{1}}{\tau_{2}}$, is negligible with respect to the second term. Thus, for a typical protein, changes in initiation rate should not sizably affect the amplitude of the observed noise, CV. In our experiments, this condition is fulfilled because the leucine CTG starvation reporter uses a ribosome binding sites whose strength is within the range of the well-characterized strong viral T7 RBS^6^. Indeed, we observe that for every leucine concentration tested, the noise is insensitive to several fold changes in initiation rate and remains within a tight band between 0.2 and 0.3 CV units, **Supplementary** **Figure 7**. This independence on translation initiation rate and its associated noise (CV) has been previously reported for cells growing in rich media^18^. Here, we observe that the dependence is also true for our reporter system in media in which amino acid (leucine), at different concentrations, is the growth limiting component.

**Strains, experimental protocol and replicates.** Using degenerate primers, we mutated the RBS region of the YFP, robust codon variant “CTG” (a member of the set of the leucine codon YFP variants) and selected seven different RBS mutants with different initiation rates. We inserted the RBS variants at the AttB site of the leucine auxotroph background strain (Noise bg V1.0, ΔleuB), see **Construction of Auxotrophs and YFP Synonymous Codon Variants**. The protocol to obtain the [leucine] response of the RBS variants was the same as that described in **Section 2.5** for the leucine set of codons. We repeated the [leucine] response of the RBS variants for another set of independent strains, i.e. a second set of colonies of successfully integrated RBS variants picked from the original transformation plates, see **Supplementary** **Figure 7**.

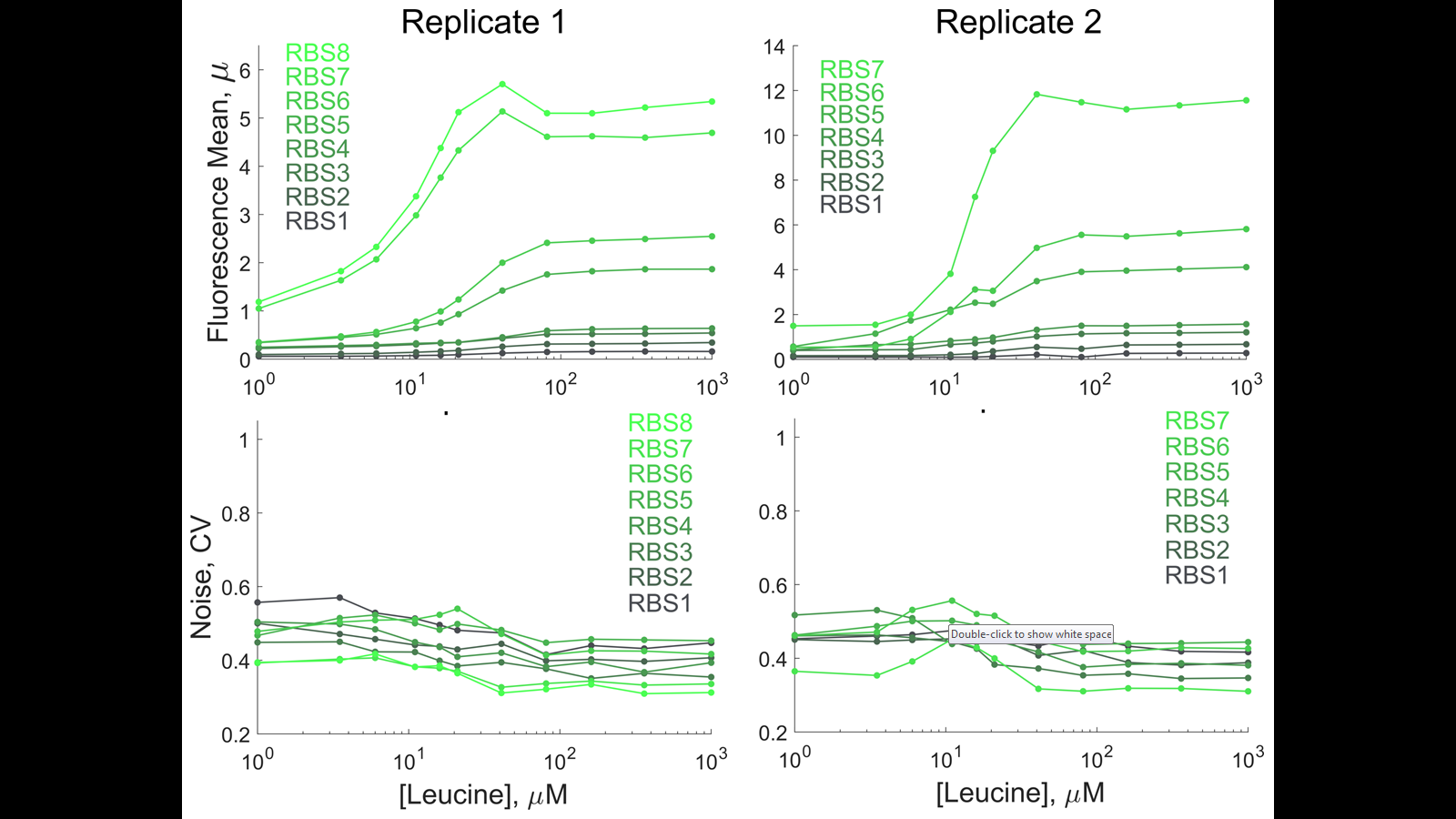

**Supplementary** **Figure 7** | **Changes** **in translation initiation rate hardly change the associated protein noise even at growth-limiting leucine concentrations.** Mean fluorescence (top) and associated noise (bottom) of a set of YFP reporters as a function of [leucine] and RBS strength. We derived all reporters from the YFP starvation reporter CTG (a robust codon from the leucine codon family) and we inserted them at the AttB site of strain Noise bg V1.0, Δ*leuB*, a strict auxotroph in synthetic media without leucine. Shades of green indicate RBS strength, from weak (black) to strong (light green). We derived mean and CV values from a sample size of 3x10^3^ ± 200 cells. Error bars associated with the standard error of the mean are smaller than symbols sizes. We performed the [leucine] response experiment twice for two sets (two bioreplicates) of RBS strains.

### 5 Two-Color Experiment

#### 5.1 2-Color Plasmid Construction, 2-Color Strains & Experimental Protocol

We cloned in a plasmid, as a bicistronic message, fast-maturing yellow (mVenNB) and cyan (SCFP3A) fluorescent proteins^20^. In order to increase signal strength in the cyan channel of the flow cytometer, we expressed the bicistronic message from plasmids instead of inserting both colors into the chromosome. The RNA message was bicistronic in order to use a single P_LtetO_^5^ promoter per plasmid—thus preventing a lack of TetR repressors due to excess of competing TetR binding sites. We cloned the construct into the low-copy SC101 origin of replication^21^. The bicistronic cassette has the CFP at its 5’ end and the YFP at its 3’ end; the same RBS (Variant#1, see **Section 1.2**) controls the translation initiation rate of each fluorescent protein. We constructed a plasmid that consists of CFP-robust & YFP-robust (robust CTG leucine codon on both colors) and another plasmid with a CFP-sensitive & YFP-sensitive (sensitive CTA leucine codon on both colors).

For the 2-color experiment, we substituted in the leucine auxotroph base strain the original cyan FP marker with a red FP marker (mCherry2). In this way, the cyan channel became available to detect the signal from the robust (or sensitive) cyan SCFP3A in the 2-color plasmid.

The protocol to obtain the [leucine] response from the 2-color plasmids was the same as that described in **Section 2.5** for the leucine set of codons.

#### 5.2 Data Analysis

To decompose local (intrinsic) and global (extrinsic) noise in the presence of systematic cell cycle variations, we follow Hilfinger and Paulsson^12^. They recognize that global noise can be overshadowed by cell cycle variations and that these variations can be taken into account by conditioning on cell size (SSC in our case). Using the law of total covariance, we can write,

$$Cov\left( CFP, YFP|SSC \right)=\underset{\begin{aligned} Global noise \\ unexplained by SSC \end{aligned}}{\underbrace{\left\langle Cov(CFP,YFP|SSC) \right\rangle}}\underset{\begin{aligned} Global noise \\ explained by SSC \end{aligned}}{+\underbrace{Cov(\left\langle CFP,YFP|SSC \right\rangle)}}$$

Thus, to obtain the global noise, $\eta_{global}$, we calculated $\eta_{global}^{2}=\left\langle Cov\left( CFP,YFP | SSC \right) \right\rangle/\langle YFP\rangle\langle CFP\rangle$. Local noise, $\eta_{local}$, is exempt from the contribution of the cell cycle variations (see equation 9 in^12^), thus to calculate it, we used the definition given in Elowitz *et al.*^22^,

$$\eta_{local}^{2}=\frac{\left\langle\left( CFP-YFP \right)^{2} \right\rangle}{2\left\langle YFP \right\rangle\left\langle CFP \right\rangle}$$

#### 5.3 Robust-Robust Data and Sensitive-Sensitive Replicate

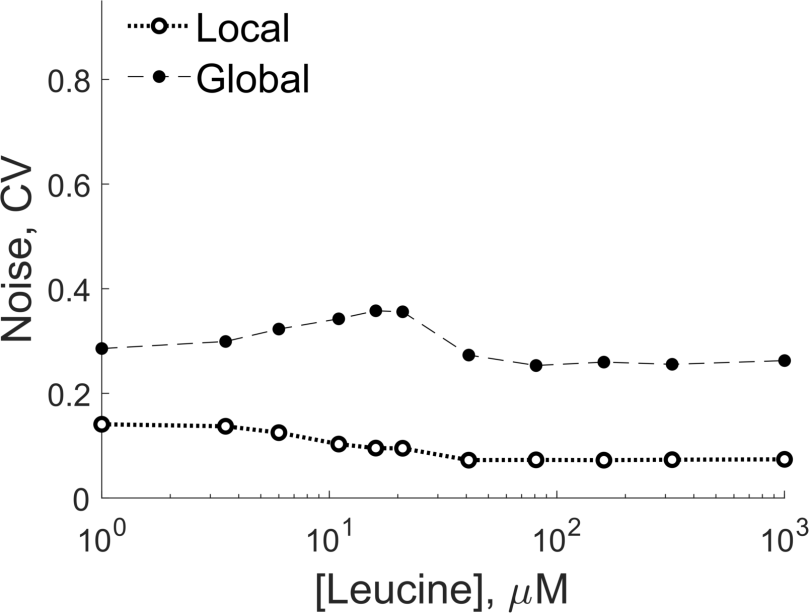

**Supplementary Figure 8** | **Noise decomposition into local and global components as a function of leucine concentration when both proteins are coded with the robust leucine codon (CTG)**. The local and global CV values were derived from a sample size of 5x10^3^ ± 200 cells. Error bars associated with the standard error of the mean are smaller than symbols sizes.

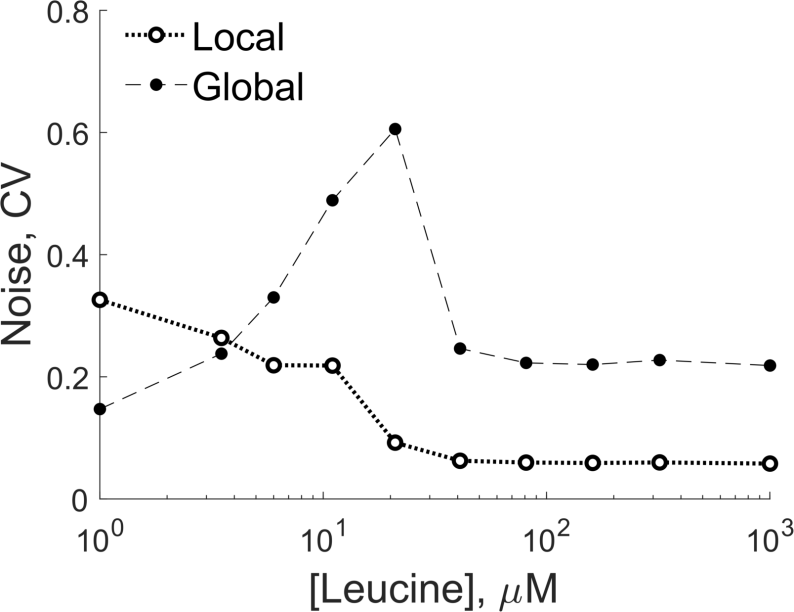

**Supplementary Figure 9** | **Noise decomposition into local and global components as a function of leucine concentration when both proteins are coded with the sensitive leucine codon (CTA); repeat of experiment in Figure 3.** The local and global CV values were derived from a sample size of 5x10^3^ ± 200 cells. Error bars associated with the standard error of the mean are smaller than symbols sizes.

### 6 tRNA Titration Experiment

#### 6.1 tRNA Strains Construction & Experimental Protocol

To titrate the amount of the leuU tRNA, tRNA^Leu^_GAG_, we constructed plasmids to drive the expression of a DNA region that included the *leuU* gene (region delimited by sequences CCGCGGCCGCATGACGGCGCTGCTGG & GGGCCCGTTGACACAATAAAGTGCC and defined by Sörensen *et al.*^23^). We varied the expression levels of *leuU* using four constitutive synthetic promoters (proA<pro5<proB<proC) that spanned a 9-fold expression range from the weakest to the strongest promoter^24^. To avoid the confounding regulated expression of the native *leuU* copy, we deleted it from the chromosome of our leucine auxotrophic background strain (Noise bg V1.0, ΔleuB). Because, *leuU* is an essential gene (it is the only tRNA that reads the CTC codon), we first cloned a ccdB cassette next to *leuU* in the chromosome. Then, we co-transformed one of the *leuU* bearing plasmids together with a linear fragment to delete the native *leuU* copy and the ccdB cassette. The linear fragment fused the first 36bp of the *leuU* gene with a downstream sequence about 270bp from the promoter of the *yhbX* gene. The linear fragment for the deletion was TACCAGCTACGAGTAAAGCAACTGGACGAGTCAAGGTAGCGTGTCTACCAATTCCACCAC.

The protocol to obtain the [leucine] response of these different tRNA strain was the same as that described in **Section 2.5** for the leucine set of codons.

### 7 tRNA Charging/Discharging Model

#### 7.1 The tRNA Charging/Discharging Model

tRNA charging is a balance between (i) an aminoacylation, $J_{S}$, reaction that charges the tRNA with its cognate amino acid and (ii) a protein synthesis reaction, $J_{R}$, mediated by a ribosome at the cognate codon, which effectively discharges tRNA. Given the typical characteristic times of the two reactions compared to the *E. coli* generation time, the total amount of tRNA can be assumed constant. A simple scheme of this charging/discharging cycle is

| 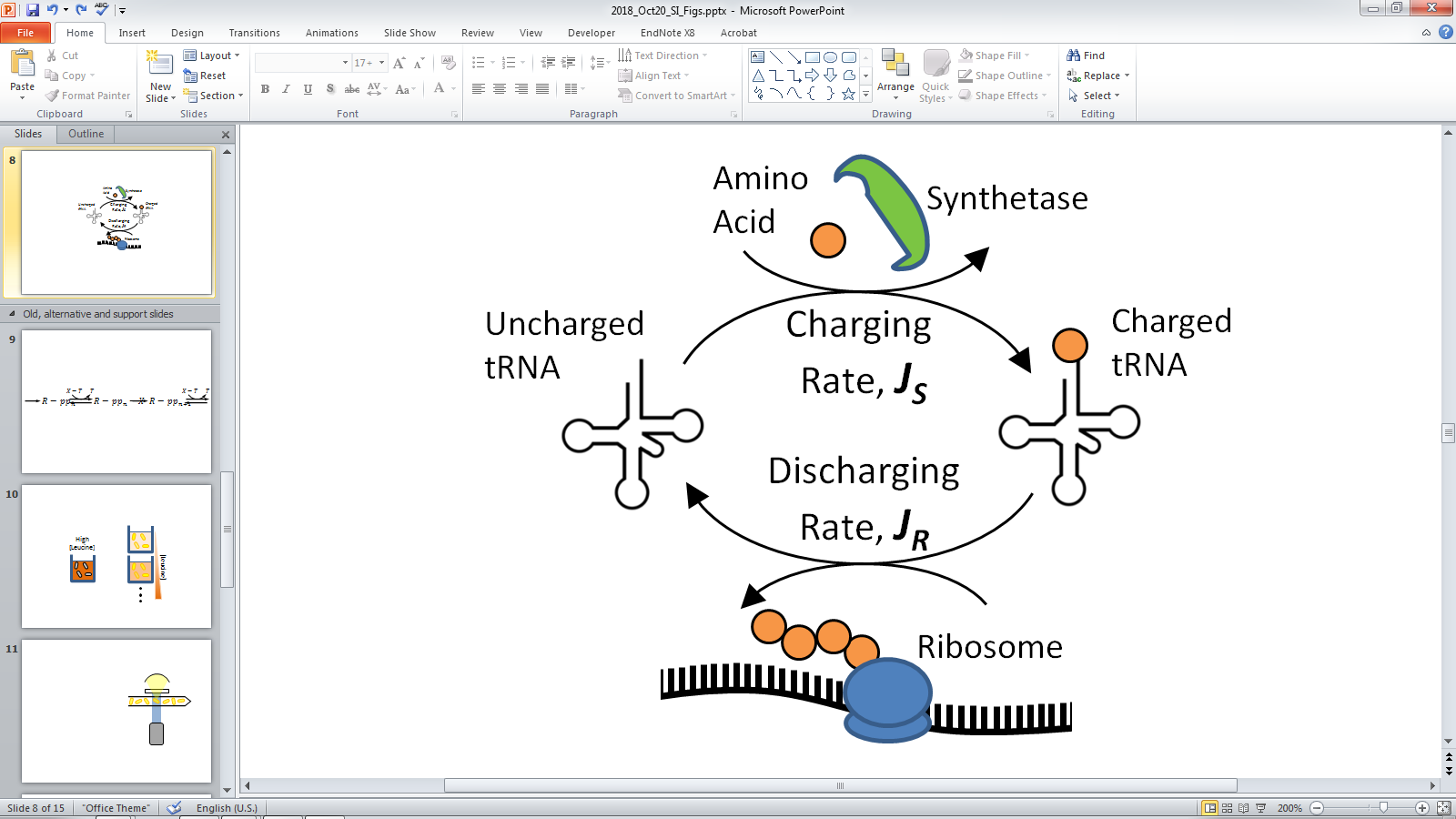 | (5) |
| --- | --- |

To model the tRNA charging and discharging reactions, we use a model proposed by Elf and Ehrenberg and published in^25-28^. The reaction scheme for tRNA charging is

| 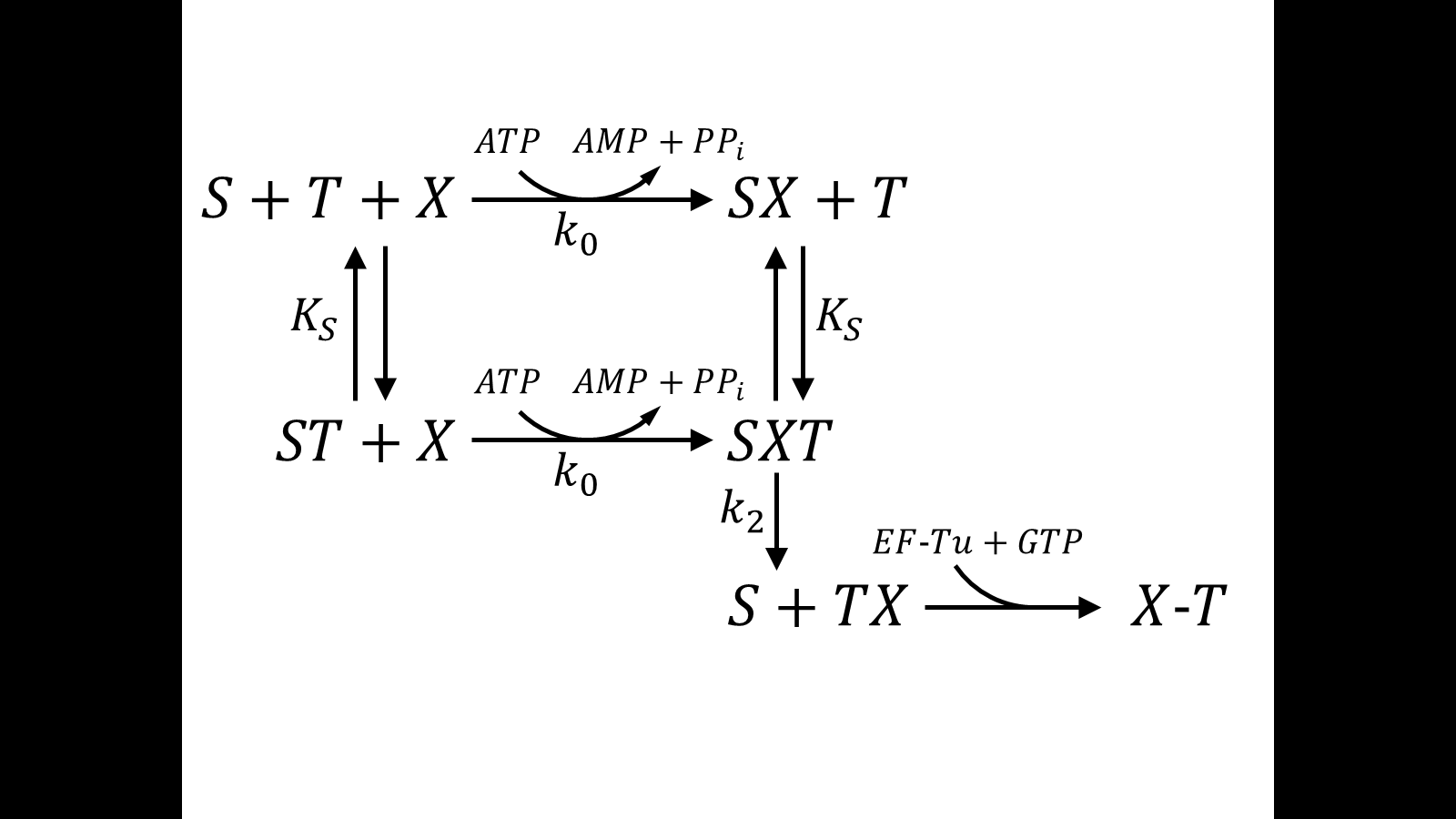 | (6) |
| --- | --- |

Where $S$ is the synthetase, $T$ the tRNA and $X$ the cognate amino acid. Independently of tRNA, the synthetase activates the amino acid with rate $k_{0}$ by using ATP and producing aminoacyl-adenylate and releasing pyrophosphate, $PP_{i}$. The tRNA binds reversibly to the synthetase with rate $K_{s}$. $k_{2}$ is the maximal turnover rate when the synthetase is saturated with tRNA and amino acid. $X$-$T$ represents the aminoacylated-tRNA in ternary complex with EF-Tu and GTP, and $SX$, $SXT$, and $ST$ represent complexes of the synthetase with amino acid, with amino acid and tRNA, and with tRNA respectively. Assuming that ATP saturates the synthetase and assuming that the tRNA equilibrates rapidly with the synthetase, the rate of tRNA aminoacylation, $J_{S}$, is

$$J_{S}=\frac{X.S{.k}_{2}}{\frac{k_{2}}{k_{0}}+X\left( 1+\frac{K_{s}}{\left( 1-\alpha\right)T} \right)}$$

(7)

Where $X$ is the internal amino acid concentration, $T$ is the tRNA concentration and $\alpha$ is the tRNA charged fraction.

The reaction scheme for tRNA discharging is

| 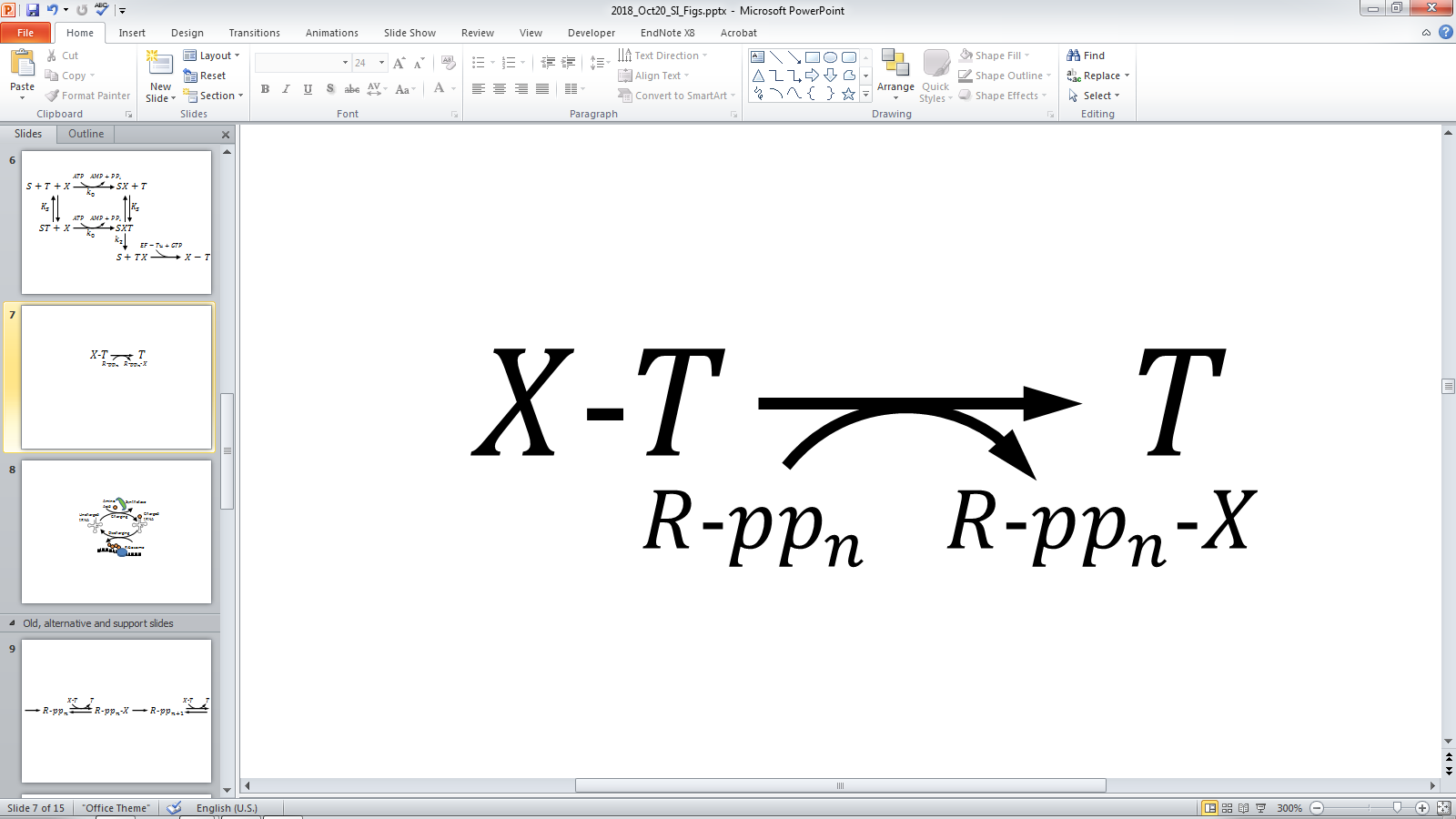 | (8) |
| --- | --- |

Where $pp_{n}$ is a nascent polypeptide bound to a ribosome $R$. The polypeptide increases irreversibly by one unit when the ribosome incorporates an amino acid bound to a tRNA. Assuming that all codons except one are translated at maximal speed, and assuming that the interaction of ribosome, charged tRNA and cognate codon can be represented by roughly the same Michaelis-Menten parameters, the rate of charged tRNA consumption by protein synthesis, $J_{R}$, is

$$J_{R}=fRv=fR\frac{k_{max}}{1+\frac{fK_{R}}{\alpha T}}$$

(9)

Where $k_{max}$ is the maximum translation rate, $K_{R}$ is the Michaelis constant Km to half-saturate with charged tRNA a translating ribosome,$v$ is the average translation speed, and $f$ is the frequency of occurrence of the limiting amino acid in proteins.

The value of the parameters are:

$k_{0}$ (M^-1^ s^-1^),	1x10^6^

$k_{2}$ (s^-1^),	100

$S$ (M),	1.3x10^-6^

$K_{s}$ (M),	1x10^-6^

$R$ (M),	2x10^-5^

$f$,	0.05

$k_{max}$ (s^-1^),	10

$K_{R}$ (M),	1x10^-6^

Reaction schemes (6) & (8) and the associated rate equations (7) & (9) can be found in^25-27^, parameter values are from^25^.

When the supply of charged tRNA equals the demand by elongating ribosomes, *i.e.* $J_{S}=J_{R}$, we can express the charged ratio $\alpha$ as a function of amino acid $X$. This is the function that we plotted in Figure 4b of main text. To generate the different curves we multiplied the wild type level of leucine tRNA by the factors: 0.25, 0.42, 1 and 2.34. The factors follow the relative changes in the *pro* series promoters^24^ proA, pro5, proB and proC. The wild type level is the total concentration of leucine tRNA which is equal to 8,775 molecules per *E. coli* volume (1 um^3^). The number of tRNA molecules is the result of adding all different tRNA species that read a leucine codon: leuPQVT= 6000, leuU = 750, leuW = 750, leuX=525, leuZ=750. The charging/discharging cycle works at saturation at as can be seen in Supplementary Fig. XX.

From the fluxes $J_{S}$ and $J_{R}$, we can calculate the response of the charged fraction, $\alpha$, to changes in amino acid, $X$, *i.e.* we can calculate the sensitivity amplification

$$a_{\alpha x}=\frac{X}{\alpha} \frac{d\alpha}{dX}=\frac{dlog(\alpha)}{dlog(X)}$$

(10)

For this calculation, we follow appendix A4 in^29^ so that

$$a_{\alpha X}=\frac{X}{\bar{\alpha}} \left( \frac{\frac{\partial J_{S}}{\partial X}}{\frac{\partial J_{R}}{\partial\alpha}-\frac{\partial J_{S}}{\partial\alpha}} \right)_{\bar{\alpha}}$$

(11)

Where $\bar{\alpha}$ is the steady state charged fraction obtained from the constrain $J_{S}=J_{R}$. We used equation (11) to plot the sensitivity shown in the inset of Fig. 4 of main text. To estimate sensitivities from experimental data, we used equation (10) and approximated the differentials by finite differences.

Note that the charging/discharging model that we are using is a minimal model in which we assume that there is only one type of tRNA that reads one type of codon. In reality, there are several types of tRNAs that are charged with the same amino acid. Moreover, a synonymous codon cannot be read by all types of tRNAs. Thus, even for a single amino acid, the translation process is degenerate. The degeneracy can be treated by using several $\alpha$’s—one for every tRNA type—and by using several$f$’s—one for every synonymous codon^28^.

#### 7.2 Noise in the tRNA Charging/Discharging Model

We used the following stochastic process to analyze fluctuations in tRNA charging,

| *In*: $X\underset{\to}{{X_{0}\lambda}_{in}}X+1$ *Out*: $X\underset{\to}{{X \lambda}_{out}}X-1$  *Charge*: $T\underset{\to}{J_{S}}T+1$ *Discharge*: $T\underset{\to}{J_{R}}T-1$ | (12) |
| --- | --- |

Reactions on the bottom row model the charging/discharging cycle with only intrinsic noise, *i.e.* noise related to the inherent probabilistic nature of the charging/discharging reactions. Reactions on the top row add to the charging/discharging process extrinsic or global noise, *i.e.* noise to the parameters that modulate the rates $J_{S}$ and $J_{R}$. In this example, we added noise to the effective amino acid concentration and labeled the reactions *In* and *Out*. Although reactions *In* and *Out* can model different processes—passive amino acid diffusion through a membrane or the internal consumption of the amino acid by other means different to charging, *e.g.* amino acid catabolism—we do not stick to a specific interpretation. Rather, reactions *In* and *Out* serve to exemplify, through the addition of simple Poissonian fluctuations to a controlling parameter, the contribution of extrinsic noise to the total noise in the system. Reaction *In* assumes a constant flux, $X_{0}\lambda_{in}$, of amino acid $x$ set by the external amino acid concentration $x_{0}$. Reaction *Out* assumes that the available amino acid is eliminated in a first order fashion, *i.e.* ${X\lambda}_{out}$. The noise associated to this stochastic process is $CV^{2}=\frac{1}{\langle X\rangle}$, thus fluctuations follow Poissonian statistics.

The complete stochastic process (12) is a pseudo-bivariate process because $T$ does not affect the rates of $X$. To analyze the fluctuations in (12), we followed the linear noise approximation given in^15^, equation 1. The result is Fig. 4c of the main text. All parameters are the same to the ones used in the macroscopic discussion of $J_{S}$ and $J_{R}$. The dashed black line in Fig. 4c is the linear noise approximation of only the bottom row of the stochastic process (12), *i.e.* the black dashed line corresponds to the local or intrinsic noise of the charging/discharging cycle. We performed all algebraic manipulations with Mathematica 11.3. The software is available as Supplementary Material.

### 8 References

1	Sharan, S. K., Thomason, L. C., Kuznetsov, S. G. & Court, D. L. Recombineering: a homologous recombination-based method of genetic engineering. *Nat Protoc* **4**, 206-223, doi:10.1038/nprot.2008.227 (2009).

2	Li, X. T., Thomason, L. C., Sawitzke, J. A., Costantino, N. & Court, D. L. Positive and negative selection using the tetA-sacB cassette: recombineering and P1 transduction in Escherichia coli. *Nucleic Acids Res* **41**, e204, doi:10.1093/nar/gkt1075 (2013).

3	Kolmsee, T. & Hengge, R. Rare codons play a positive role in the expression of the stationary phase sigma factor RpoS (sigma(S)) in Escherichia coli. *RNA Biol* **8**, 913-921, doi:10.4161/rna.8.5.16265 (2011).

4	Nielsen, H. J., Li, Y., Youngren, B., Hansen, F. G. & Austin, S. Progressive segregation of the Escherichia coli chromosome. *Mol Microbiol* **61**, 383-393, doi:10.1111/j.1365-2958.2006.05245.x (2006).

5	Lutz, R. & Bujard, H. Independent and tight regulation of transcriptional units in Escherichia coli via the LacR/O, the TetR/O and AraC/I1-I2 regulatory elements. *Nucleic Acids Res* **25**, 1203-1210 (1997).

6	Olins, P. O. & Rangwala, S. H. A novel sequence element derived from bacteriophage T7 mRNA acts as an enhancer of translation of the lacZ gene in Escherichia coli. *J Biol Chem* **264**, 16973-16976 (1989).

7	Kudla, G., Murray, A. W., Tollervey, D. & Plotkin, J. B. Coding-Sequence Determinants of Gene Expression in Escherichia coli. *Science* **324**, 255-258, doi:10.1126/science.1170160 (2009).

8	Goodman, D. B., Church, G. M. & Kosuri, S. Causes and effects of N-terminal codon bias in bacterial genes. *Science* **342**, 475-479, doi:10.1126/science.1241934 (2013).

9	Baba, T. *et al.* Construction of Escherichia coli K-12 in-frame, single-gene knockout mutants: the Keio collection. *Mol Syst Biol* **2**, 2006 0008, doi:10.1038/msb4100050 (2006).

10	Flint, K. P. The long-term survival of Escherichia coli in river water. *J Appl Bacteriol* **63**, 261-270 (1987).

11	Poncelet, P. *et al.* Standardized counting of circulating platelet microparticles using currently available flow cytometers and scatter-based triggering: Forward or side scatter? *Cytometry A* **89**, 148-158, doi:10.1002/cyto.a.22685 (2016).

12	Hilfinger, A. & Paulsson, J. Separating intrinsic from extrinsic fluctuations in dynamic biological systems. *Proc Natl Acad Sci U S A* **108**, 12167-12172, doi:10.1073/pnas.1018832108 (2011).

13	Nordholt, N., van Heerden, J., Kort, R. & Bruggeman, F. J. Effects of growth rate and promoter activity on single-cell protein expression. *Sci Rep* **7**, 6299, doi:10.1038/s41598-017-05871-3 (2017).

14	Bagwell, C. B., Hill, B. L., Herbert, D. J., Bray, C. M. & Hunsberger, B. C. Sometimes simpler is better: VLog, a general but easy-to-implement log-like transform for cytometry. *Cytometry A* **89**, 1097-1105, doi:10.1002/cyto.a.23017 (2016).

15	Paulsson, J. Summing up the noise in gene networks. *Nature* **427**, 415-418, doi:10.1038/nature02257 (2004).

16	Elf, J. & Ehrenberg, M. Fast evaluation of fluctuations in biochemical networks with the linear noise approximation. *Genome Res* **13**, 2475-2484, doi:10.1101/gr.1196503 (2003).

17	Bar-Even, A. *et al.* Noise in protein expression scales with natural protein abundance. *Nat Genet* **38**, 636-643, doi:10.1038/ng1807 (2006).

18	Ozbudak, E. M., Thattai, M., Kurtser, I., Grossman, A. D. & van Oudenaarden, A. Regulation of noise in the expression of a single gene. *Nat Genet* **31**, 69-73, doi:10.1038/ng869 (2002).

19	Milo, R. & Phillips, R. *Cell biology by the numbers*. (Garland Science, Taylor & Francis Group, 2016).

20	Balleza, E., Kim, J. M. & Cluzel, P. Systematic characterization of maturation time of fluorescent proteins in living cells. *Nat Methods* **15**, 47-51, doi:10.1038/nmeth.4509 (2018).

21	Cox, R. S., 3rd, Dunlop, M. J. & Elowitz, M. B. A synthetic three-color scaffold for monitoring genetic regulation and noise. *J Biol Eng* **4**, 10, doi:10.1186/1754-1611-4-10 (2010).

22	Elowitz, M. B., Levine, A. J., Siggia, E. D. & Swain, P. S. Stochastic gene expression in a single cell. *Science* **297**, 1183-1186, doi:10.1126/science.1070919 (2002).

23	Sorensen, M. A. *et al.* Over expression of a tRNA(Leu) isoacceptor changes charging pattern of leucine tRNAs and reveals new codon reading. *J Mol Biol* **354**, 16-24, doi:10.1016/j.jmb.2005.08.076 (2005).

24	Davis, J. H., Rubin, A. J. & Sauer, R. T. Design, construction and characterization of a set of insulated bacterial promoters. *Nucleic Acids Res* **39**, 1131-1141, doi:10.1093/nar/gkq810 (2011).

25	Elf, J., Berg, O. G. & Ehrenberg, M. Comparison of repressor and transcriptional attenuator systems for control of amino acid biosynthetic operons. *J Mol Biol* **313**, 941-954, doi:10.1006/jmbi.2001.5096 (2001).

26	Elf, J. *Intracellular Flows and Fluctuations* PhD thesis, Uppsala Universitet, (2004).

27	Elf, J. & Ehrenberg, M. Near-critical behavior of aminoacyl-tRNA pools in E. coli at rate-limiting supply of amino acids. *Biophys J* **88**, 132-146, doi:10.1529/biophysj.104.051383 (2005).

28	Elf, J., Nilsson, D., Tenson, T. & Ehrenberg, M. Selective charging of tRNA isoacceptors explains patterns of codon usage. *Science* **300**, 1718-1722, doi:10.1126/science.1083811 (2003).

29	Elf, J., Paulsson, J., Berg, O. G. & Ehrenberg, M. Near-critical phenomena in intracellular metabolite pools. *Biophys J* **84**, 154-170, doi:10.1016/S0006-3495(03)74839-5 (2003).
